# Immunolocalization studies of vimentin and ACE2 on the surface of cells exposed to SARS-CoV-2 Spike proteins

**DOI:** 10.1101/2021.05.04.442648

**Authors:** Vasiliki Lalioti, Silvia González-Sanz, Irene Lois-Bermejo, Patricia González-Jiménez, Álvaro Viedma-Poyatos, Andrea Merino, María A. Pajares, Dolores Pérez-Sala

## Abstract

The Spike protein from SARS-CoV-2 mediates docking of the virus onto cells and contributes to viral invasion. Several cellular receptors are involved in SARS-CoV-2 Spike docking at the cell surface, including ACE2 and neuropilin. The intermediate filament protein vimentin has been reported to be present at the surface of certain cells and act as a co-receptor for several viruses; furthermore, its potential involvement in interactions with Spike proteins has been proposed. Here we have explored the binding of Spike protein constructs to several cell types using low-temperature immunofluorescence approaches in live cells, to minimize internalization. Incubation of cells with tagged Spike S or Spike S1 subunit led to discrete dotted patterns at the cell surface, which showed scarce colocalization with a lipid raft marker, but consistent coincidence with ACE2. Under our conditions, vimentin immunoreactivity appeared as spots or patches unevenly distributed at the surface of diverse cell types. Remarkably, several observations including potential antibody internalization and adherence to cells of vimentin-positive structures present in the extracellular medium exposed the complexity of vimentin cell surface immunoreactivity, which requires careful assessment. Notably, overall colocalization of Spike and vimentin signals markedly varied with the cell type and the immunodetection sequence. In turn, vimentin-positive spots moderately colocalized with ACE2; however, a particular enrichment was detected at elongated structures positive for acetylated tubulin, consistent with primary cilia, which also showed Spike binding. Thus, these results suggest that vimentin-ACE2 interaction could occur at selective locations near the cell surface, including ciliated structures, which can act as platforms for SARS-CoV-2 docking.

## Introduction

Vimentin is an intermediate filament cytoskeletal protein highly abundant in mesenchymal cells. In addition, several other cell types express vimentin under normal circumstances, including endothelial cells, certain cells lining the airways and several cells of the immune system (https://www.proteinatlas.org/ENSG00000026025-VIM). Moreover, vimentin expression can be increased or induced in cells under situations of injury, senescence or epithelial-mesenchymal transition, as it occurs during tumorigenesis or chronic inflammation [1-3]. Vimentin plays important mechanical functions in cells providing support, sustaining the position and protection of cellular organelles, including the nucleus [4-7], and interacting with the other cytoskeletal systems to contribute to essential cellular functions such as division and migration [8-10]. In addition to its functions in the intracellular environment, vimentin has been reported to exert important actions at the cell surface or in the extracellular medium (reviewed in [11, 12]). Vimentin has been detected at the surface of diverse cell types where it acts as a receptor for several types of ligands, such as soluble CD44 [13] or certain carbohydrate chains [14]. Moreover, vimentin has been reported to function as a ligand for various receptors, including IGF-1R [15] and P-selectin [16]. Circulating forms of vimentin, either in extracellular vesicles or in other non-completely characterized forms, have been described [17-19], which could be involved in autoimmunity, epithelial mesenchymal transition or in the modulation of inflammatory responses. Importantly, extracellular and/or cell surface vimentin species have been reported to act as co-receptors for several pathogens, including bacteria and viruses, either facilitating cellular invasion or acting as a restriction factor for this process [20, 21] (reviewed in [12, 22]). In the context of viral invasion, vimentin was reported to act as a co-receptor for SARS-CoV, the coronavirus causing the 2003 outbreak, in association with the protein ACE2 [23]. The need for therapeutic tools in the fight against SARS-CoV-2, the virus causing the COVID-19 pandemic, has fostered the interest in vimentin as a potential therapeutic target in viral infections, a subject addressed in several recent reviews and hypotheses [12, 24, 25].

Research on SARS-CoV-2 represents one of the most extraordinary efforts in Biomedicine, and an enormous amount of resources have been devoted to understand and combat this new pathogen in a record time. The study of SARS-CoV-2 entails high complexity due to the variety of syndromes and symptoms it can provoke. Initially thought to produce a flu-like disease, it is now clear that this virus can invade multiple cell types in the organism and trigger, directly or indirectly, dysfunction of almost any system, and provoke the severe cytokine storm that contributes to the high mortality of this disease [26, 27]. In an *in vitro* assay platform SARS-CoV-2 was able to enter cardiomyocytes, pancreatic beta cells, liver organoids and dopaminergic neurons [28]. Indeed, human small intestine organoids are also infected by the virus and support viral replication [29].

Spike proteins are glycoproteins present at the outer layer of the virus that mediate binding of the viral particles to cellular receptors and membrane fusion. The best characterized receptor for SARS-CoV-2 is ACE2, and detailed structural information on the Spike-ACE2 interaction is available [30, 31]. Nevertheless, the ability of this virus to enter diverse cell types [28] could be related to its capacity to interact with other cellular receptors, as it has been reported for integrins [32], neuropilin [33], and the tyrosine-protein kinase receptor UFO (AXL) [34]. In addition, membrane-bound lectins, heparan sulfates or specific lipid domains could favor the interaction of the virus with the cell surface [35-38]. The Spike protein possesses two domains which are generated by posttranslational cleavage by furin, but remain associated thereafter. The S1 domain is involved in the interaction with ACE2 through the so-called receptor binding domain (RBD), whereas the S2 domain is implicated in membrane fusion [31]. Interestingly, both, viral particles and recombinant Spike chimeric proteins containing either the full length protein or specific domains, i.e., S1 or RBD, have been used to explore the interaction with cellular receptors in cell-based assays, as well as *in vitro* [39, 40]. Here, we have imaged different cell types upon incubation with several recombinant versions of the Spike protein.

Moreover, we have attempted the detection of vimentin and/or ACE2 on the cell surface before and after incubation with the Spike protein constructs. Our observations highlight the complexity of vimentin immunodetection at the cell surface. Moreover, a highly consistent Spike-ACE2 colocalization, but low ACE2-vimentin and Spike-vimentin coincidence were observed. Nevertheless, the three proteins appear to concur at certain cellular structures, in particular, primary cilia, the importance of which in Spike docking deserves further investigation.

## Materials and Methods

Recombinant proteins. SARS-CoV-2 Spike protein constructs, Spike S1-Fc-Avi and Spike S1-His-Avi (containing residues Gln14 to Arg683 of the Spike protein), Spike S-Fc-Avi (S1+S2; containing amino acids Gln14 to Trp1212), and human ACE2-His-Avi (residues Gln18 to Ser740), all expressed in HEK293 cells, were from Bioss Antibodies. Human IgG1-Fc protein “103Cys/Ser” was from Sino Biological. Cholera toxin subunit B-Alexa555 (CTXB) was from Molecular Probes.

Antibodies. Anti-vimentin antibodies used were: anti-vimentin antibody V9 (sc-6260, unconjugated and Alexa-488-conjugate) from Santa Cruz Biotechnology; mouse monoclonal anti-vimentin clone 13.2 (V5255) from Sigma; rabbit monoclonal SP20 anti-vimentin antibody from ThermoFisher Scientific; cell-surface vimentin (CSV) antibody, clone 84-1, from Abnova, and goat anti-vimentin antibody EB11207 from Everest Biotech. Anti-ARL13B C-5 (sc-515784) was from Santa Cruz Biotechnology, anti-ACE2 rabbit antibodies were purchased from Invitrogen and Abcam, anti-human IgG-Alexa647 and anti-human IgG-Alexa568 were obtained from Invitrogen, anti-acetylated tubulin (T7451) was from Sigma and anti-SARS-CoV-2 Spike protein was a product of Sino Biological. Details of the antibodies used are summarized in Supplementary Table 1.

Cell culture. Cell culture media and supplements were from Gibco. Fetal bovine serum (FBS) was from Sigma. Fibroblast-like African green monkey kidney cells (Vero) were from the collection of Centro de Investigaciones Biológicas Margarita Salas (Madrid). Human adrenal carcinoma SW13/cl.2 cells were the generous gift of Dr. A. Sarriá (University of Zaragoza, Spain) [41]. SW13/cl.2 cells stably expressing vimentin wt and RFP as separate products (RFP//vimentin wt) have been previously described [6, 9]. Cells were cultured in DMEM supplemented with 10% (v/v) FBS and antibiotics (100 U/ml penicillin, 100 μg/ml streptomycin). Stably transfected cell lines were maintained in the presence of 500 μg/ml geneticin. Human lung adenocarcinoma A549 cells (ATCC) were cultured in RPMI1640 with 10% (v/v) FBS, 50 U/ml penicillin, 50 μg/ml streptomycin and 50 μg/ml gentamycin. HAP1 cells are near haploid human cells derived from a chronic myelogenous leukemia cell line, which present an adherent epithelioid-like phenotype. HAP1 cells, parental (vimentin positive, vim +), and vimentin-depleted (vim -) through CRISPR-CAS9-engineering, (reference number HZGHC003297c010) were from Horizon. HAP1 cells were cultured in DMEM-F12 with 10% (v/v) FBS, 100 U/ml penicillin, 100 μg/ml streptomycin, and 2 mM glutamine (Gibco).

Immunofluorescence studies. Cells were grown on glass coverslips. In the basic immunodetection protocol, coverslips were washed with cold PBS and transferred to a plate cover lined with Parafilm placed on ice. Cells were incubated with 10 μg/ml Spike protein constructs (final concentration) in 1% (w/v) BSA in PBS for 1 h. Unless specified otherwise, antibodies were routinely used at 1:200 (v/v) dilution in 1% (w/v) BSA in PBS, and incubations were carried out for 1 h. For incubations, coverslips were kept over an ice bed and at 4ºC, and washes were performed on ice with cold PBS. Controls of all secondary antibodies, alone and in combination, as well as others omitting particular primary or secondary antibodies, were performed to ensure specificity of the signals detected. When a particular reagent was omitted, cells were incubated with 1% (w/v) BSA in PBS for an equivalent time. Whenever possible, observations were confirmed using different primary antibodies against the same protein or tag. For labeling with CTXB-Alexa555 conjugate, cells on coverslips were incubated with 0.25 μg/ml CTXB in 1% (w/v) BSA in PBS for 10 min on ice, either at the beginning or at the end of the procedure involving incubation with Spike constructs plus immunodetection, and subsequently washed with cold PBS. Finally, cells were fixed with 4% (w/v) paraformaldehyde (PFA) for 25 min on ice. After washing with PBS and water, coverslips were allowed to dry at r.t. and mounted onto glass slides using Fluorsave (Calbiochem). Where indicated, cells were fixed with 4% (w/v) PFA for 25 min on ice before vimentin immunodetection. In addition, several alternative protocols were used for detection of intracellular proteins or markers of primary cilia. In some assays, cells were fixed with 4% or 2% (w/v) PFA for 15 min at r.t., subsequently incubated with 0.1% (v/v) Triton X-100 for 5 min, 50 mM glycine for 20 min, and blocked with 1% (w/v) BSA in PBS for 1 h before immunodetection with anti-acetylated tubulin at 1:800 dilution, or anti-vimentin and anti-ARL13B at 1:400 dilution, and the corresponding secondary antibodies at 1:250 dilution, in both cases through incubation for 1 h at 37ºC. Nuclei were stained with 4,6-diamidino-2-phenylindole (DAPI) from Sigma, at 3 μg/ml in PBS. Variations in these procedures are specified in the figure legends.

Fluorescent labeling of Spike S1-Fc. A 10 μl aliquot of Spike S1 at 1 mg/ml as provided by the manufacturer was incubated for 20 min at r.t in the presence of a 10-fold molar excess of FITC or a 7-molar excess of Alexa Fluor 488 carboxylic acid succinimidyl ester (CASE) dye (ThermoFisher Scientific). Afterwards, excess reagent was removed by filtration through a Zeba desalting micro-spin column (ThermoFisher Scientific). The filtrate was stored at -80ºC until used.

Confocal microscopy and image analysis. Images were acquired on SP5 or SP8 Leica confocal microscopes, using 63x or 100x oil immersion objectives. Single confocal sections were taken every 0.5 μm in sequential mode. Images were analyzed using LasX or ImageJ software. For analysis of colocalization of the signals of interest at the plasma membrane, band-like regions of interest (ROIs) extending 2 µm outwards from the internal side of the plasma membrane were defined for individual cells in single z-sections where the highest membrane-located signal was observed. Colocalization was analyzed using Fiji’s JACoP plugin and was presented as percentages of coincidental signals (Mander’s coefficient), under the threshold determined by Costes’s method. Colocalization masks were obtained with LasX software.

Cell lysis and western blot. Cell monolayers were washed with ice-cold PBS. For analysis of vimentin, HAP1 and SW13/cl.2 cells were homogenized in 50 mM Tris-HCl pH 7.5, 0.1 mM EDTA, 0.1 mM EGTA, 0.1 mM β-mercaptoethanol, 0.5% (w/v) SDS, 20 mM sodium orthovanadate, 50 mM sodium fluoride, containing protease inhibitors (2 μg/ml each of leupeptin, aprotinin and trypsin inhibitor, and 1.3 mM Pefablock). For assessment of ACE2 levels, A549 and Vero cells were lysed in RIPA containing cOmplete™ Protease Inhibitor Cocktail (Sigma). Cell debris was removed by centrifugation at 16000xg for 5 min at 4ºC. Protein concentration was determined by the Bicinchoninic acid method (Pierce, ThermoFisher Scientific Fisher, Rockford, IL, USA). Aliquots of lysates containing 30 μg of protein were separated on SDS-PAGE and transferred to Immobilon-P membranes, essentially as described [42]. After blocking with 2% (w/v) non-fat dried milk, blots were incubated with primary antibodies, typically at 1:500 dilution, followed by HRP-conjugated secondary antibodies, at 1:2000 dilution. In all cases, polypeptides of interest were visualized with the enhanced chemiluminiscence system (ECL, GE Healthcare). Total protein on blots was stained with Simply Blue Colloidal Coomassie reagent (Invitrogen).

Statistical analysis. Statistical analyses were performed with GraphPad Prism 5 software. Results are shown as mean values ± standard error of the mean (SEM). The colocalization extent upon the different immunofluorescence incubation protocols was statistically compared, for every cell type, by ANOVA with Tukey’s post-test. Non-significant (ns, p > 0.05) and significant (*, p ≤ 0.05; **, p ≤ 0.01; ***, p ≤ 0.001) differences are indicated in the graphs.

## Results

### Detection of Spike protein constructs at the cell surface

In order to detect Spike bound to the cell surface, Vero cells were incubated with Spike S1-Fc protein and subjected to immunofluorescence. All the procedure was carried out on ice to avoid internalization of the viral protein. This resulted in a dotted pattern on the cell surface clearly detectable by confocal microscopy using Alexa-conjugated anti-human IgG or anti-Fc antibodies (Fig. 1A). Incubation with a Spike S-Fc construct, bearing the S1 and S2 domains, yielded a similar pattern on the cell surface (Fig. 1A). In all cases, the anti-IgG or anti-Fc conjugates gave only background staining. Moreover, incubation of cells with the IgG1-Fc fragment resulted in negligible staining, indicating that the cellular binding of Spike S1-Fc protein constructs is not mediated by the Fc tag (Fig. 1A). Cell-bound tagged Spike S1 proteins could also be visualized with an anti-Spike antibody, which gave a clean background in cells incubated with vehicle (Fig. 1B). Finally, a fluorescent Spike S1-Fc protein labeled with CASE-Alexa488 retained the ability to bind to cells yielding a dotted pattern on the cell surface (Fig. 1C). Moreover, overlapping of the signal of Spike S1-Fc-Alexa488 and that obtained by detection of the Fc tag with Alexa647-anti-human IgG confirmed the specificity of the detection (Fig. 1C).

**Figure 1.**
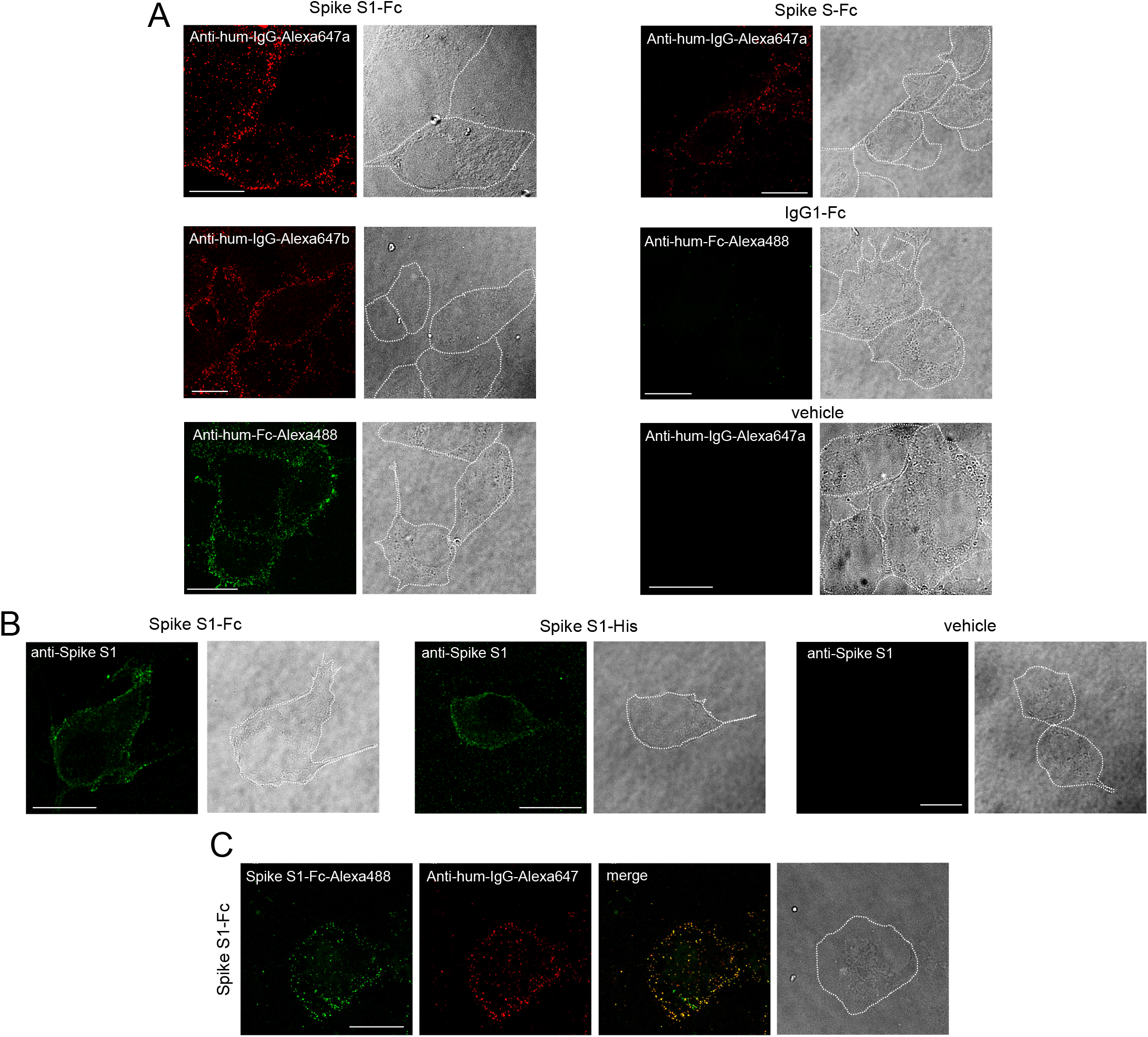
Detection of Spike protein constructs at the surface of Vero cells. (A) Cells plated on glass coverslips were incubated with recombinant Spike S1-Fc, Spike S-Fc, Fc or vehicle, as indicated, for 1 h in the cold, as described in Methods, and the proteins retained on the cell surface were detected with the indicated antibodies. Anti-human (anti-hum) IgG-Alexa647a and Anti-human-IgG-Alexa647b are antibodies raised in goat and alpaca, respectively. (B) Vero cells were incubated as above with Spike S1-Fc, Spike S1-His or vehicle, as indicated, and Spike constructs were detected with an anti-Spike S1 antibody. (C) Cells were incubated with Spike S1-Fc labeled with Alexa488 (green) followed by anti-human IgG-Alexa647 (red), and the overlay of the two signals is shown (merge). In all cases, bright field images in which the cell contours have been highlighted are shown. Bars, 20 μm.

Binding of Spike proteins to the cell surface could also be evidenced in other fibroblast-like cells, namely, human HAP-1 cells, and in epithelial cells such as adrenal carcinoma SW13/cl.2 and lung adenocarcinoma A549 cells (Fig. 2), the latter reported to express variable levels of ACE2 protein [43] but lacking Fc receptors [44]. Importantly, Spike S and S1 protein constructs could be detected after incubation for several hours at 4ºC and extensive washing, indicating high binding stability. Nevertheless, a variable degree of signal internalization even at this low temperature was observed in several images. Moreover, the distribution of Spike constructs was not totally homogeneous in the cell population, with some cells or cell areas showing more intense staining, as illustrated for instance for Vero cells (Fig. 2, arrows).

**Figure 2.**
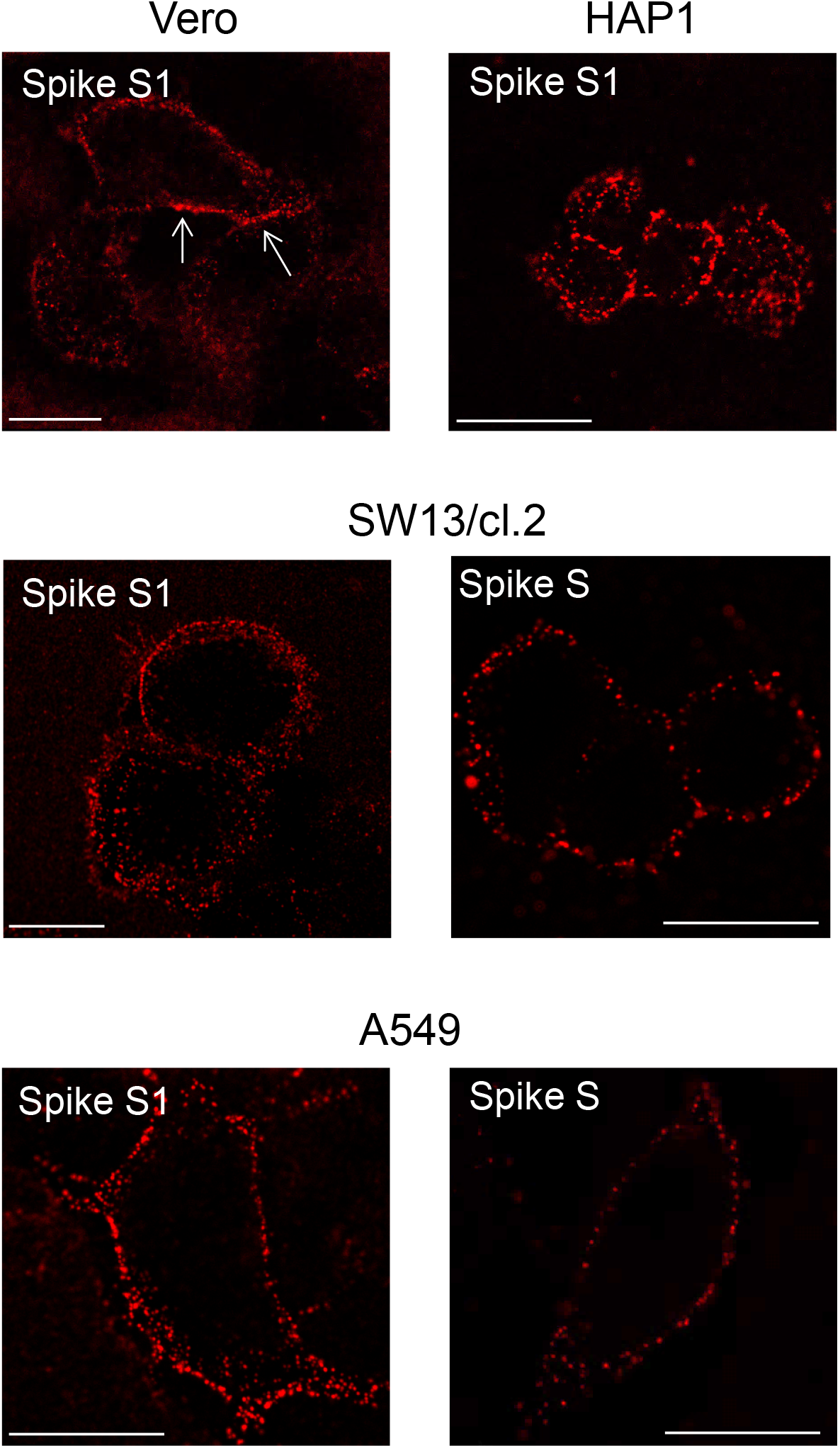
Detection of Spike protein constructs in several cell types. Fibroblast-like Vero and HAP1 cells, or epithelial SW13/cl.2 and A549 cells, were incubated with Spike S-Fc (S) or Spike S1-Fc (S1), as above, and detected by incubation with anti-human-IgG-Alexa647. Arrows point to regions with more intense Spike signal. Bars, 20 μm.

### Spike protein constructs display scarce colocalization with cholera toxin B-positive membrane domains

Several studies have proposed the importance of lipid rafts in SARS-CoV-2 infection. Therefore, as an additional parameter to characterize the binding of Spike constructs to the cell surface, we stained cells with fluorescent cholera toxin B subunit (CTXB), which binds the lipid membrane component GM1 ganglioside, and is widely used as a marker for membrane microdomains with the characteristics of lipid rafts [45]. Labeling with CTXB was also carried out at 4ºC in order to minimize endocytosis of the toxin. CTXB-Alexa555 stained regions or discrete patches of the membrane (Fig. 3), which showed scarce colocalization with the Spike S1-Fc signal (Fig. 3A), as indicated by a Mander’s colocalization coefficient of 0.007 ± 0.002 (mean ± SEM of 42 measurements, under the threshold determined by the Costes’ method). Spike fluorescent spots showed some partial overlapping with CTXB-positive patches, as evidenced in the fluorescence intensity profiles (Fig. 3A). As an example, 3 out of 13 Spike S1-Fc-positive peaks coincided with CTXB fluorescence peaks in the representative profile shown in Fig. 3A. As CTXB has the property to bind up to five molecules of its lipid receptor, thus, associating with and crosslinking lipid rafts, it may potentially remodel the underlying membrane [45]. To discard potential effects of CTXB *per se*, labeling with the toxin was also performed after incubating cells with Spike-Fc protein constructs (Fig.3B). We observed that the calculated colocalization extent was also virtually negligible (0 ± 0.0002, n=14), with some Spike S1-Fc-positive peaks coinciding with CTXB fluorescence peaks in the illustrative profile shown in Fig. 3B. In addition, some apparently intracellular patches of CTXB could be detected, which also showed scarce colocalization with Spike S1-Fc. Therefore, under our conditions, binding of Spike S1 to the cell surface did not appear to occur specifically at lipid rafts.

**Figure 3.**
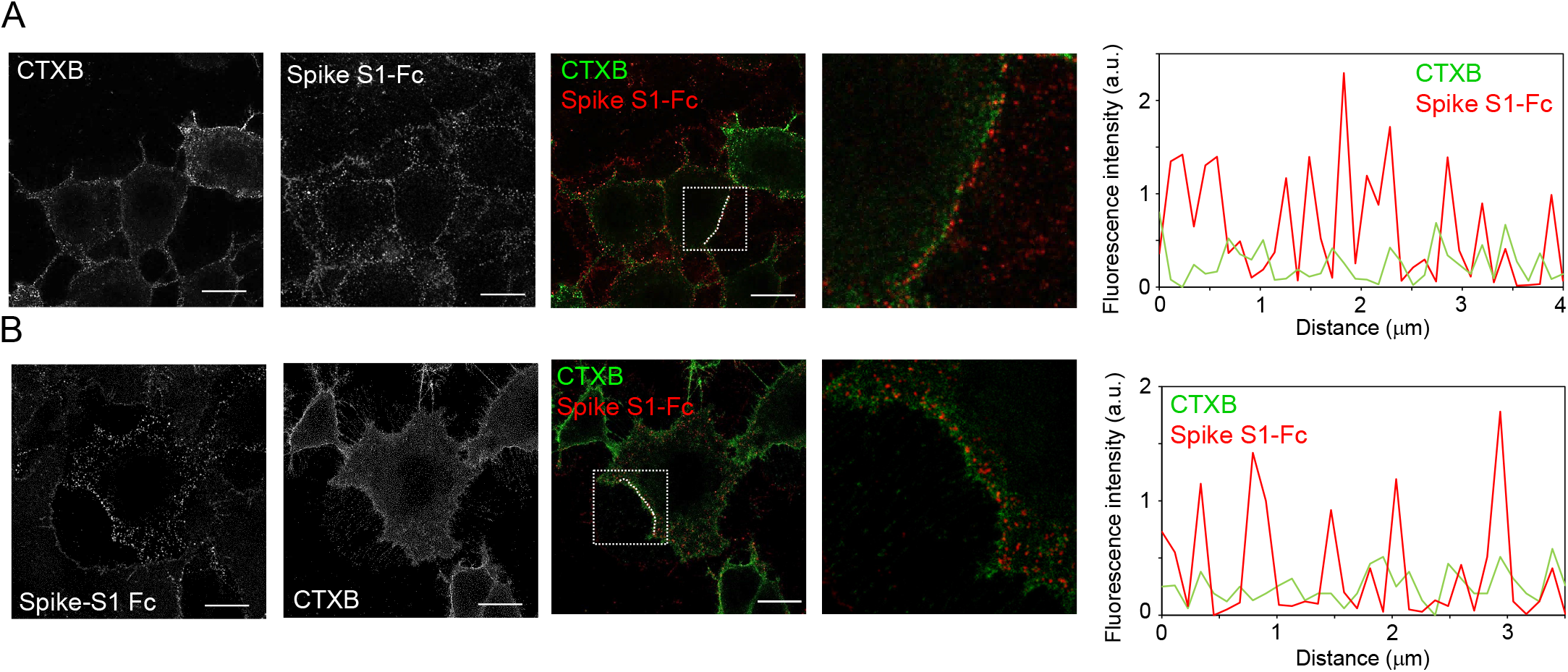
Detection of Spike S1 and cholera toxin-B binding sites in cells. Vero cells on glass coverslips were incubated in the cold with cholera toxin-B--Alexa555 (CTXB) for 10 min before (A) or after (B) incubation with Spike S1-Fc for 1 h followed by anti-human IgG-Alexa647 for another hour. Cells were extensively washed after each incubation. At the end of the procedure, cells were fixed and mounted for visualization by confocal microscopy. Images of individual channels and a merged image of a section at mid-cell height are shown. Regions of interest in the merged images, delimited by dotted squares, are magnified on the right. Graphs on the far right display the fluorescence intensity profiles of Spike S1-Fc (red) and CTXB (green) signals along the dotted lines marked in the merged images. Bars, 20 μm.

### Detection of vimentin at the cell surface

The exposure of vimentin at the cell surface has been reported in several cell types in association with cell senescence and/or oxidative damage [46] (reviewed in [12]). Nevertheless, the disposition of the protein at the cell surface is not well understood, and the epitopes exposed appear to vary with the experimental model used [23, 47]. Here we have employed a panel of anti-vimentin antibodies, most of which are monoclonal antibodies, some of them tested in knockout cells (“knockout validated”), as detailed in Supplementary Table 1, to assess the presence of vimentin immunoreactive signals at the cell surface.

Vero cells incubated with antibodies against vimentin showed a variable degree of staining that generally appeared as a punctate pattern at the cell surface (Fig. 4A). This staining was not homogeneous in the cell population, but was distinct from the background signal obtained with the secondary antibody (Fig. 4A). Of the antibodies used, clone 84-1 gave consistent signals under the same experimental conditions. Interestingly, the 84-1 antibody has been reported to recognize “cell-surface vimentin”, or CSV, in several cell types. Moreover, 84-1 immunoreactivity is considered as a marker for tumor cells [48]. The 84-1 epitope has not been unequivocally characterized, but epitope mapping studies are consistent with the preferential recognition of a sequence compatible with the peptide ^106^ELQELNDR^113^ (numbering from Met1) [49], located at the beginning of the rod domain. Other monoclonal antibodies, namely, V9, which recognizes the vimentin C-terminal “tail” domain, and clone 13.2, which recognizes a non-characterized epitope outside the tail domain [9], gave very faint signals. Similar results were obtained in the A549 lung carcinoma cell line, which also showed clear binding of the 84-1 antibody at the cell periphery, although it was more intense in certain cells or cell areas. In contrast, peripheral immunoreactivity with the V9 or the 13.2 antibodies was much weaker and dispersed (Fig. 4A). In addition, we tested a rabbit monoclonal anti-vimentin antibody of undefined epitope (SP20), which gave a nearly negligible signal at the cell periphery in non-permeabilized Vero or A549 cells fixed under the same conditions; however, this antibody yielded a peripheral dotted signal upon fixing with a lower concentration of PFA (2% (w/v))(Suppl. Fig. 1). Similarly, a goat polyclonal antibody recognizing the C-terminal end of vimentin also yielded a punctate peripheral pattern after 2% (w/v) PFA fixation in A549 cells (Suppl. Fig. 1). Adequate performance of the antibodies was confirmed by assessing their ability to recognize filamentous vimentin by conventional immunofluorescence protocols employing fixed and permeabilized cells (Fig. 4B and Suppl. Fig. 1). This ruled out that the faint detection of extracellular vimentin could be due to poor antibody performance.

**Figure 4.**
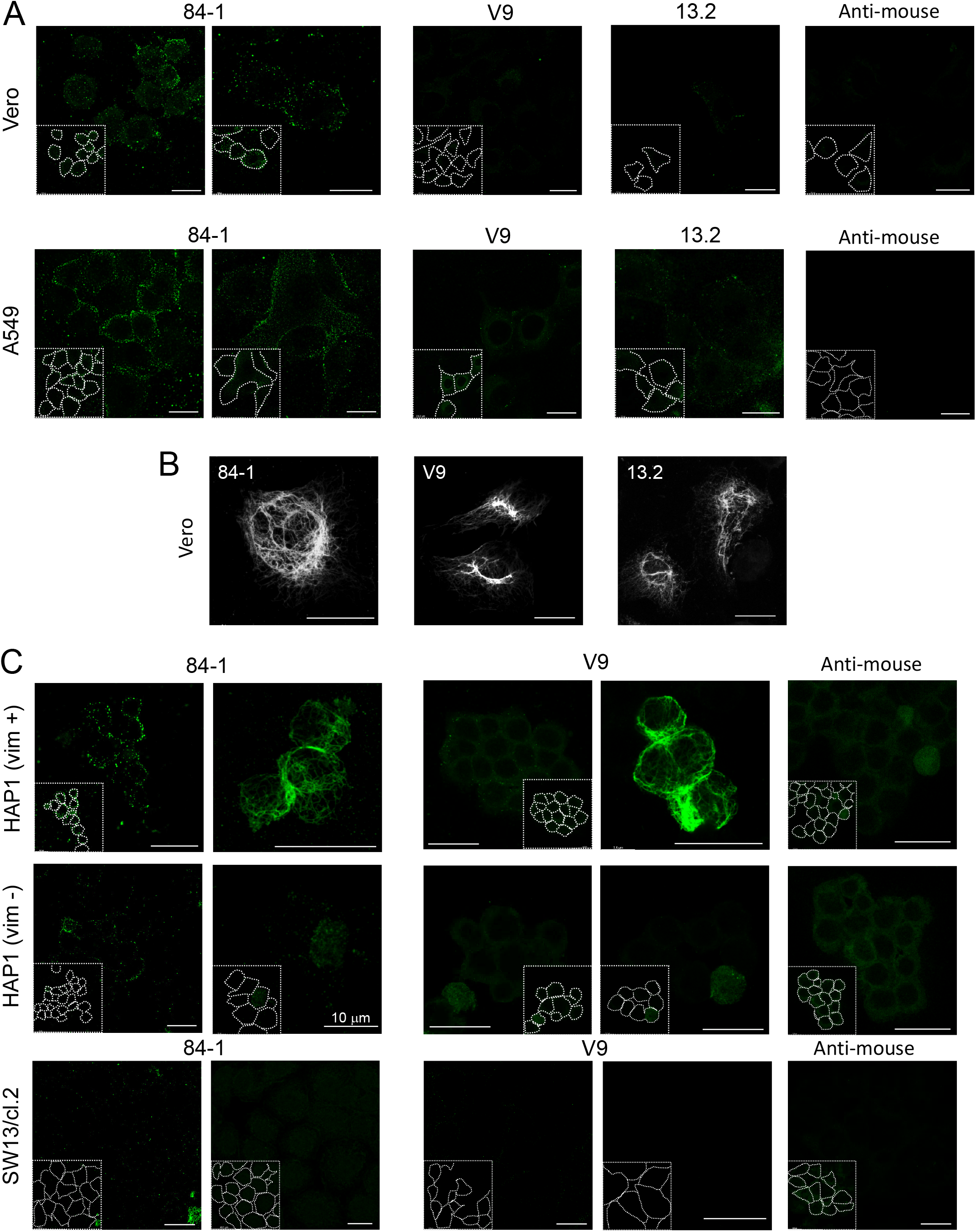
Detection of vimentin in several cell types. (A) Vero or A549 cells were incubated with the indicated monoclonal primary antibodies at 1:200 dilution for 1 h in the cold, followed by incubation with Alexa-488-conjugated anti-mouse immunoglobulins at 1:200. At the end of the procedure cells were fixed with 4% (w/v) PFA in the cold. Representative images at mid-cell height are shown. Illustrative images of the negligible background signal of secondary antibodies in every cell line are shown in the far right panels. Two representative images are depicted for the 84-1 antibody. (B) Vero cells were fixed and permeabilized for detection of cytoplasmic vimentin with the same monoclonal antibodies used in A. (C) Vimentin-positive HAP(vim +), vimentin-negative HAP(vim -), or vimentin-deficient SW13/cl.2 cells, were incubated with the indicated anti-vimentin monoclonal antibodies and Alexa-488-conjugated anti-mouse immunoglobulins at 1:200, before fixation (left images) or after fixation and permeabilization (right images). Insets display the approximate cell contours drawn from bright field or overexposed images. Images on the far right show the background of the secondary antibody in these cell types. Bars, 20 μm, unless otherwise indicated.

To ensure the specificity of detection, 84-1 and V9 antibodies were used on vimentin-positive and negative cells. HAP1 parental cells (vim +) presented a robust 84-1 immunoreactive signal in the form of bright dots or small aggregates at the cell periphery, displaying variable intensity and distribution (Fig. 4C). In HAP1 vimentin knockout cells (vim -), 84-1 gave a low, although non-negligible punctate background. The faint peripheral signal obtained with V9 was virtually undetectable in HAP1 (vim -) cells. After permeabilization, both antibodies detected filamentous vimentin in HAP (vim +) cells, whereas vimentin filaments were totally absent from HAP1 (vim -) cells. Some scattered cells with a weak intracellular dotted pattern were observed, which could represent partially damaged cells. In addition, we explored 84-1 and V9 immunoreactivity in the vimentin-deficient cell line SW13/cl.2 (Fig. 4C). In these cells, both antibodies gave a faint irregular dotted background. The secondary antibody gave a negligible or diffuse background in the various cell lines. Neither the V9 [9] nor the 84-1 antibody (Suppl. Fig. 2) detected any band in total lysates from vimentin-deficient cells by western blot.

Therefore, several anti-vimentin antibodies yielded a punctate pattern at the cell surface, which was more defined with the 84-1 antibody, denoting certain specificity, although the nature of the species detected requires further characterization.

### Additional phenomena leading to vimentin detection at the cell surface

Notably, in most of the preparations of Vero or A549 cells there was a small proportion of cells that showed staining of small areas of cytoplasmic vimentin, frequently under the plasma membrane, as illustrated in the examples presented in Fig. 5A (upper two rows). This effect could be due to focal membrane damage, occurring either spontaneously in cell culture or during the experimental procedure, which could compromise membrane integrity allowing partial internalization of the antibody. Additionally, cells can release vimentin subunits or filament fragments that can be detected in the extracellular medium. As illustrated in Fig. 5A (lower row), some filament fragments or aggregates appeared adhered to the surface of some cells. Notably, when cells were fixed with 4% (w/v) PFA before immunodetection, a substantial proportion of the cell population showed nearly complete permeabilization (Fig. 5B, lower panels), thus indicating that selective detection of cell surface vimentin is not compatible with prior fixation with this method. Hence, several phenomena related to cell damage may contribute to the presence and detection of vimentin on or at the cell surface by a variety of techniques and denote the complexity of this process. Of note, cell culture in the absence of serum did not reduce vimentin staining in A549 cells or background in vimentin-deficient SW13/cl.2 cells (Suppl Fig. 3), indicating that proteins from serum do not constitute an important source of vimentin immunoreactivity under the conditions employed.

**Figure 5.**
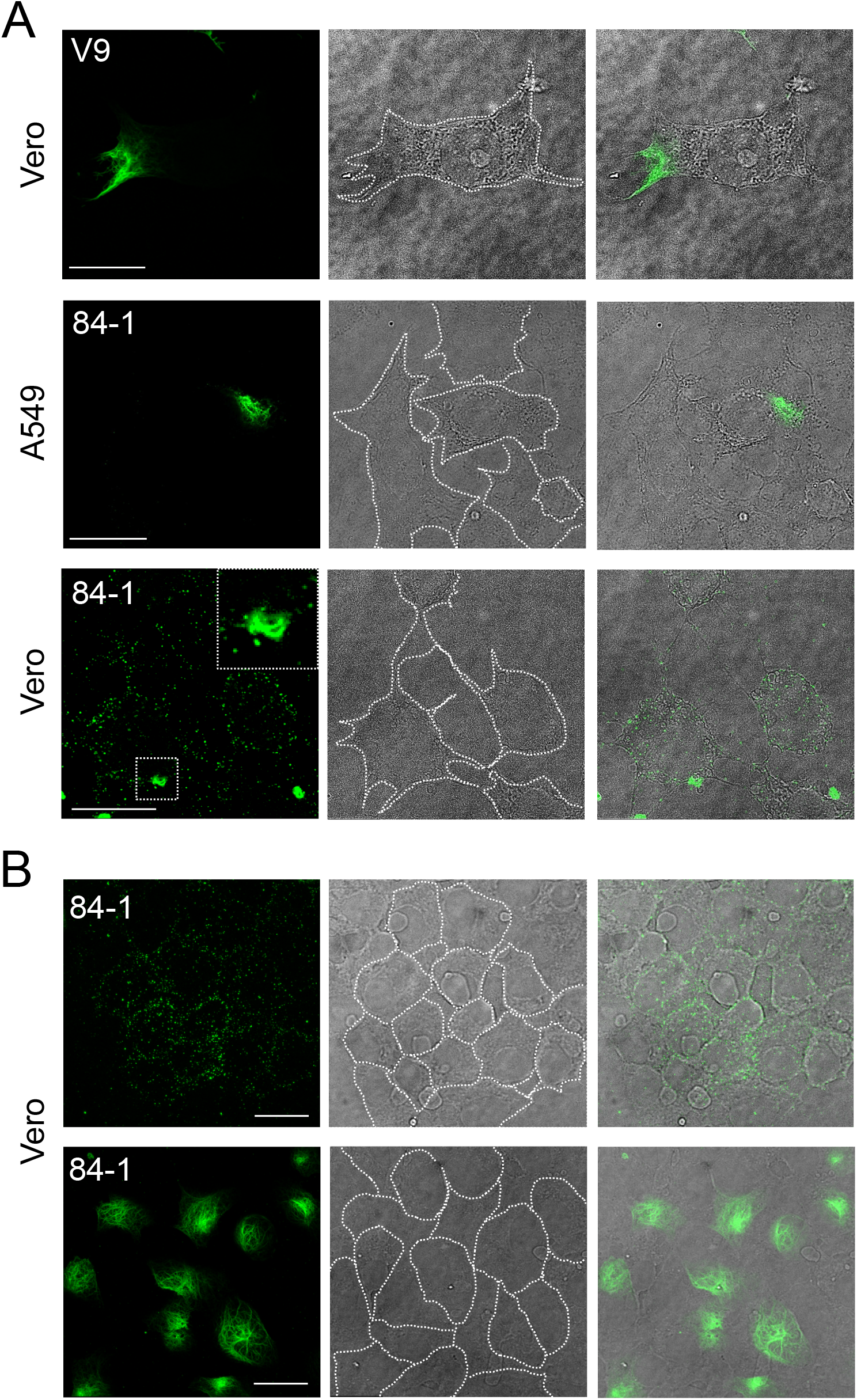
Examples of vimentin detection at or near the cell surface is cells not subjected to detergent permeabilization. (A) Vero or A549 cells were incubated with the indicated anti-vimentin antibodies as described above for detection of cell surface vimentin. Images shown illustrate representative cases of cells in which an area of the cytoplasm shows staining of filamentous vimentin (upper two rows), or that display fragments of vimentin filaments associated to their surface (lower panels). (B) Vero cells were stained with anti-vimentin antibodies prior (upper panels) or after fixation with 4% (w/v) PFA in the cold (lower panels). No additional permeabilization step was performed. Bars, 20 μm.

### Detection of ACE2

ACE2 has been extensively reported to act as a receptor for SARS-CoV-2 and to directly interact with its Spike protein both *in vitro* and in cells [37, 50]. Therefore, we attempted the detection of endogenous ACE2 in several cell types. In non-permeabilized Vero and A549 cells a polyclonal antibody against the N-terminus of the protein (anti-ACE2 p) yielded a regular peripheral staining (Fig. 6A). A polyclonal antibody against the carboxyl-terminal domain (anti-ACE2 ab) also yielded peripheral dots in non-permeabilized cells, which were clearly distinguishable from the background of the secondary antibody (Fig. 6A). At present, we do not have an explanation for this pattern since the epitope of anti-ACE2 ab reportedly corresponds to the cytoplasmic domain. In permeabilized A549 cells, anti-ACE2 p detected mainly juxtanuclear structures compatible with Golgi localization in cells cultured for 5 days after passage, but showed a more disperse cytoplasmic distribution in cells cultured for 7 days (Fig. 6B). In turn, anti-ACE2 ab showed a more diffuse pattern. Moreover, both antibodies detected several bands consistent with ACE2 in western blots of total lysates from Vero and A549 cells, although with different efficiency (Suppl. Fig. 4). These bands likely represent different posttranslationally processed forms of the protein, as it has been recently characterized [51]. Both antibodies recognized a broad 120 kDa band in Vero cells and a doublet in A549 cells. These signals coincided approximately with the position of a commercial ACE2 glycosylated protein, purified from HEK293 cells, which spans residues Gln18-Ser740 (Suppl. Fig. 4), and that could be detected by anti-ACE2 p but not by anti-ACE2 ab, consistent with the specificity of the latter antibody against the C-terminal end of the protein, as depicted in Suppl. Fig. 4C. A doublet at approximately 75 kDa, consistent with the ACE2 peptide spanning residues from 130 to 805 was recognized by both antibodies in the two cell types, whereas smaller products, likely representing cleaved forms, were preferentially recognized by the anti-ACE2 p antibody. These recognition patterns are consistent with the diverse forms of ACE2 reported in cells [51] and with the localization of the respective antibody epitopes, as schematized in Suppl. Fig 4C. Taken together, these results indicate that the anti-ACE2 p and anti-ACE2 ab antibodies detect ACE2 through various techniques although yielding different patterns, potentially related to their ability to recognize differently processed forms of the protein. Based on these data, the anti-ACE2 p antibody was preferentially used for subsequent experiments, although some results were confirmed with anti-ACE2 ab.

**Figure 6.**
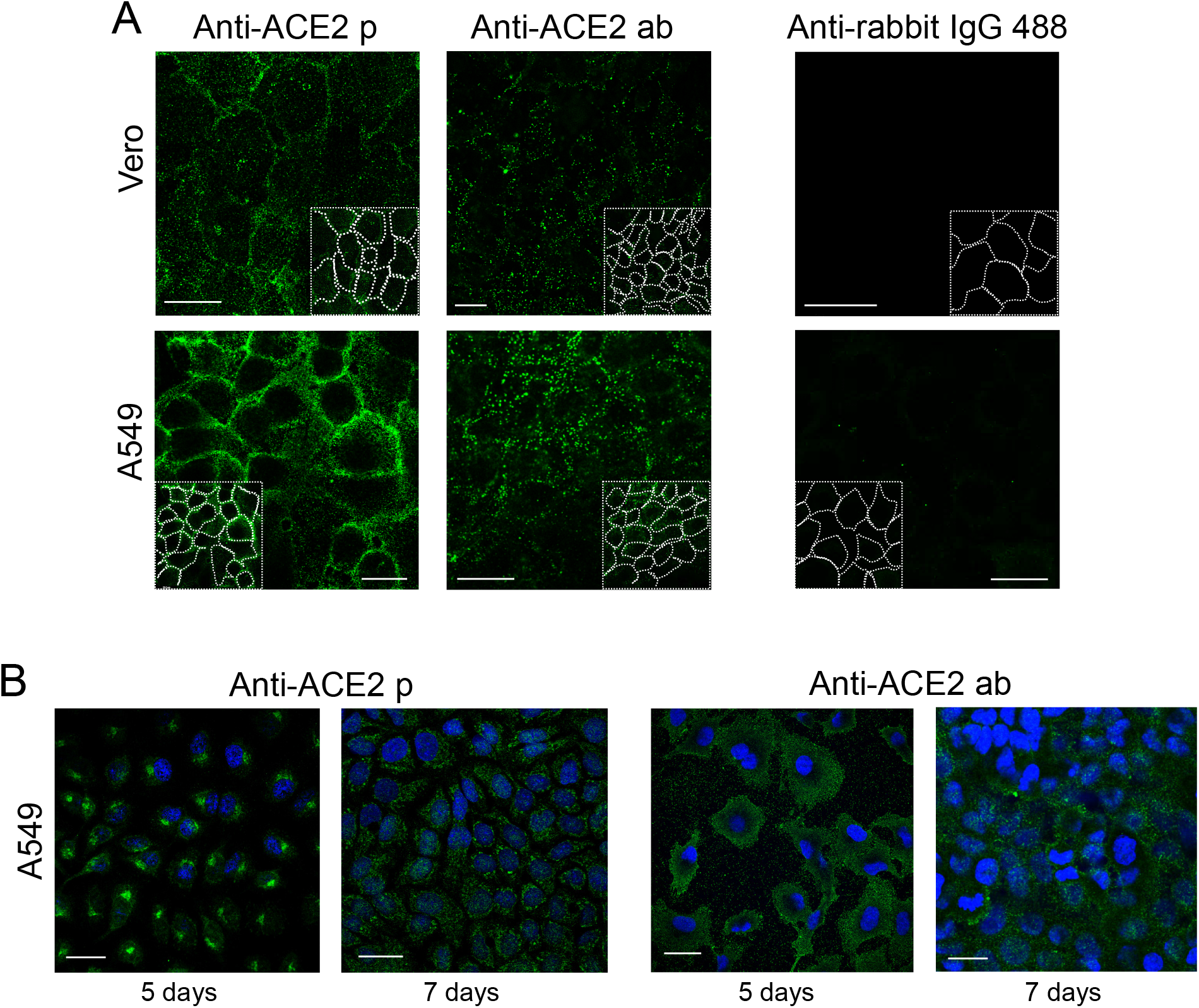
Immunodetection of ACE2. (A) Vero or A549 cells were incubated with the indicated anti-ACE2 antibodies prior to fixation. Right panels show the background of the secondary anti-rabbit IgG antibody. Insets show the cell contours. (B) A549 cells cultured for 5 days or 7 days after plating were stained with the indicated anti-ACE2 antibodies after fixation with 4% (w/v) PFA and permeabilization with 0.1% (v/v) Triton X-100 for 5 min. Nuclei were counterstained with DAPI. (p, rabbit polyclonal; ab, rabbit polyclonal antibody from Abcam). Bars, 20 μm.

### Detection of Spike protein constructs and vimentin on the surface of cells

In order to assess the relative position of Spike positive spots and vimentin immunoreactive signals at the cell surface we employed several sequential immunofluorescence protocols in live cells, schematized in Fig. 7A. Briefly, in sequence A, cells were incubated first with Spike S1-Fc and the corresponding anti-human IgG antibody prior to vimentin immunodetection, whereas in sequence B incubations were carried out in the opposite order. Finally, in sequence C, cells were incubated first with the primary reagents, i.e., Spike S1-Fc and anti-vimentin, prior to incubation with the secondary antibodies. Although several anti-vimentin antibodies were initially used, the 84-1 antibody was preferred given its better performance in terms of signal intensity and reproducibility, background and specificity in dot blot (see below). Illustrative examples of the results obtained are shown in Fig. 7B, which depicts images taken at mid-height of the cells. Manders coefficients were determined for quantitation of the percentage of Spike colocalizing with the vimentin immunoreactive signal and the results are summarized in the graph.

**Figure 7.**
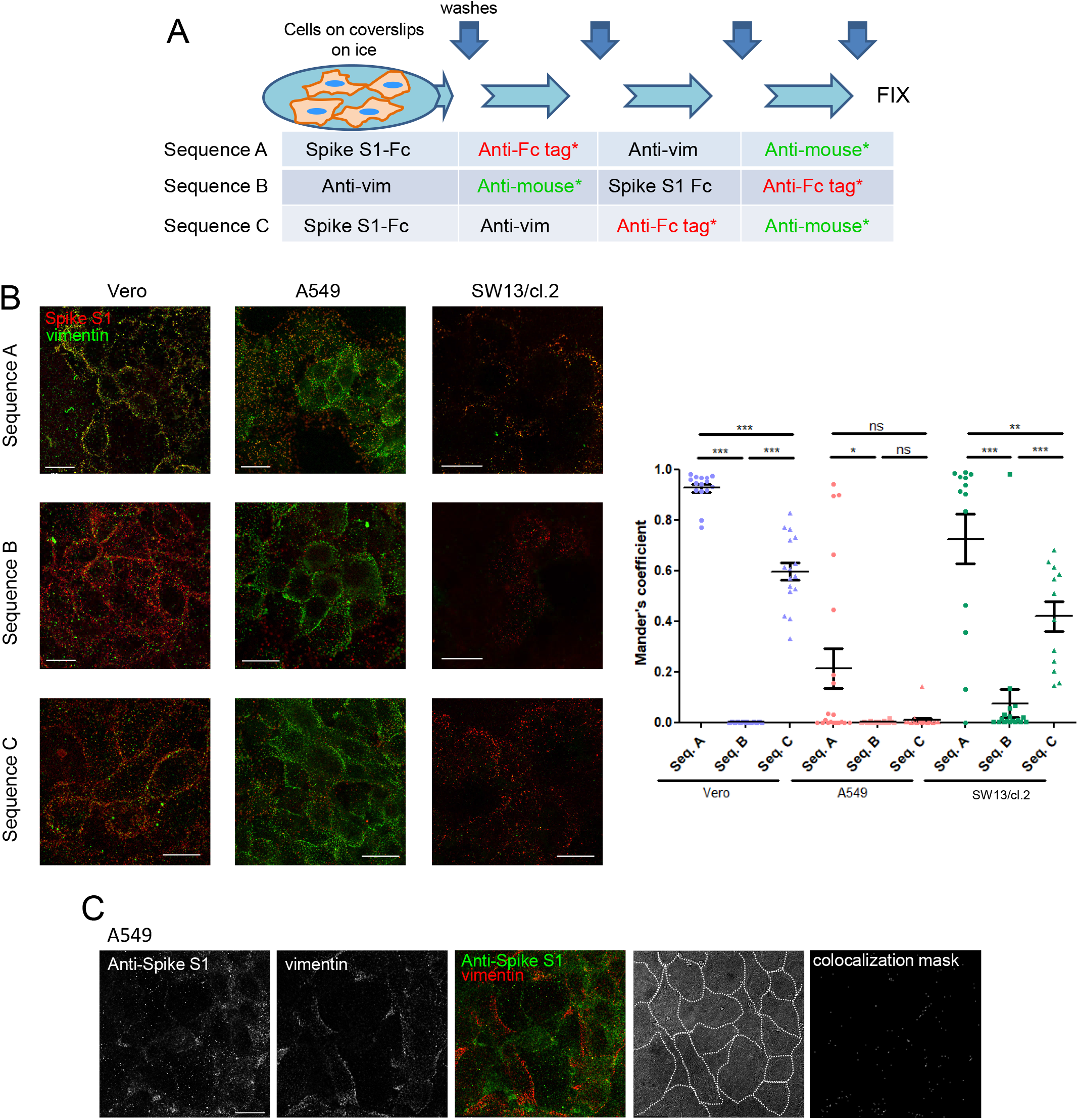
Detection of Spike S1 and vimentin in several cell types employing different immunodetection sequences. (A) Scheme of the incubation and washing steps performed for immunodetection. Incubations were carried out for 1 h in the cold. After each incubation coverslips were washed three times with 200 μl of cold PBS. At the end of the procedure, an additional washing step with water was performed before coverslips were allowed to dry and mounted. (B) Representative images from the detection of Spike S1-Fc and vimentin in the indicated cell lines following the different immunodetection sequences. The graph shows the colocalization between Spike and vimentin fluorescent signals for every cell type and immunodetection sequence assayed. Colocalization is expressed as the percentage of Spike S1 colocalizing with vimentin signal (Mander’s coefficient), measured applying the automatic Costes’ threshold (n≥12 per condition). Results are shown as mean values ± SEM; ns, non-significant, p > 0.05; *, p ≤ 0.05; **, p ≤ 0.01; ***, p ≤ 0.001 by ANOVA with Tukey’s post-test. (C) A549 cells were incubated with Spike S1-Fc, after which, immunodetection was performed by simultaneous incubation with anti-S1 antibody and 84-1 anti-vimentin monoclonal antibody, followed by simultaneous incubation with the two corresponding secondary antibodies prior to fixation. The bright field image with cell contours highlighted and colocalization mask are shown at the right. Bars, 20 μm.

In Vero cells incubated in sequence A, both Spike S1 and vimentin could be detected at the cell surface, showing a significant colocalization extent (Fig. 7B, left column). Nevertheless, when sequence B was employed, colocalization was virtually abolished. Immunodetection employing sequence C led to an intermediate staining pattern and colocalization extent. Of note, vimentin fragments adhered on the surface of cells did not show any consistent enrichment of Spike binding (Suppl. Fig. 5).

In A549 cells, the pattern of cell surface vimentin was similar independently of the strategy used for immunodetection (Fig. 7B, middle column images). Nevertheless, the extent of colocalization also varied with the immunodetection sequence. In samples processed in sequence A, a partial and widely variable coincidence of Spike and vimentin-positive spots was observed. Curiously, patches of the membrane with more intense or continuous vimentin showed a scarce presence of Spike signals. In this cell type, the extent of colocalization was also blunted by performing immunodetection according to sequence B, but did not improve significantly when sequence C was employed. Therefore, it appears that the extent of colocalization between Spike S1 and vimentin-positive signals varied markedly depending on the cell type and the sequence of the incubation steps in the detection protocol.

Next, we explored the detection of Spike S1and vimentin in vimentin-deficient SW13/cl.2 cells. As shown above in Fig. 2, Spike S1-Fc was detected at the surface of these cells, whereas the anti-vimentin antibody gave a residual signal (Fig. 7B, right column images). However, in these cells a paradoxical result was obtained since, although the signal of the anti-vimentin antibody was virtually undetectable compared to the other cell types, the calculated extent of colocalization with Spike yielded broadly variable, but unexpectedly high values in sequences A and C. Therefore, confirmation of colocalization using several antibodies and immunodetection protocols is advisable.

The higher vimentin signal and colocalization with Spike S1 observed when processing the various cell types in sequence A could be related to several factors, including potential crossreactivity between the antigens/reagents present in the incubations. To obtain additional information on a potential crossreactivity between different antibodies we performed dot blots. The results depicted in Suppl. Fig. 6, showed that the various antibodies used displayed a variable degree of crossreactivity with other immunoglobulins. Out of the primary anti-vimentin antibodies employed, 84-1 was the one showing higher specificity in terms of higher vimentin signal and lower crossreactivity, whereas V9 gave a slightly higher background with some immunoglobulins and clone 13.2 gave a low vimentin signal. Taken together, these observations indicate that, even if some of the antibodies used are highly specific, i.e., knockout validated, binding to other immune complexes or new epitopes formed during the experiment could occur, for which caution needs to be exercised in this type of experiments.

Hence, the potential colocalization of Spike S1 and vimentin at the surface of A549 cells was evaluated by an alternative protocol based on the detection with anti-Spike antibodies. The results, shown in Fig. 7C indicate a low extent of colocalization of the Spike S1 protein construct and vimentin immunoreactive signals when assessed by this strategy, as suggested by image analysis (Pearson’s correlation 0.11; overlap coefficient, 0.35; colocalization rate, 12%), and illustrated by the colocalization mask (Fig. 7C, far right image).

### Detection of Spike protein constructs and ACE2 on the surface of cells

Similar strategies to those depicted in Fig. 7A, were used to explore the colocalization of ACE2 with Spike proteins (detailed in Fig. 8A). The anti-ACE2 p antibody gave a clear signal at the cell periphery of non-permeabilized cells in all cell types tested (Fig. 8B). This pattern was similar regardless of the immunodetection sequence employed. Moreover, a high colocalization rate of Spike with ACE2 signals, measured at mid-height of the cells, was obtained in all conditions and cell types (Fig. 8B, graph), and was particularly evident at some cell-cell contacts.

**Figure 8.**
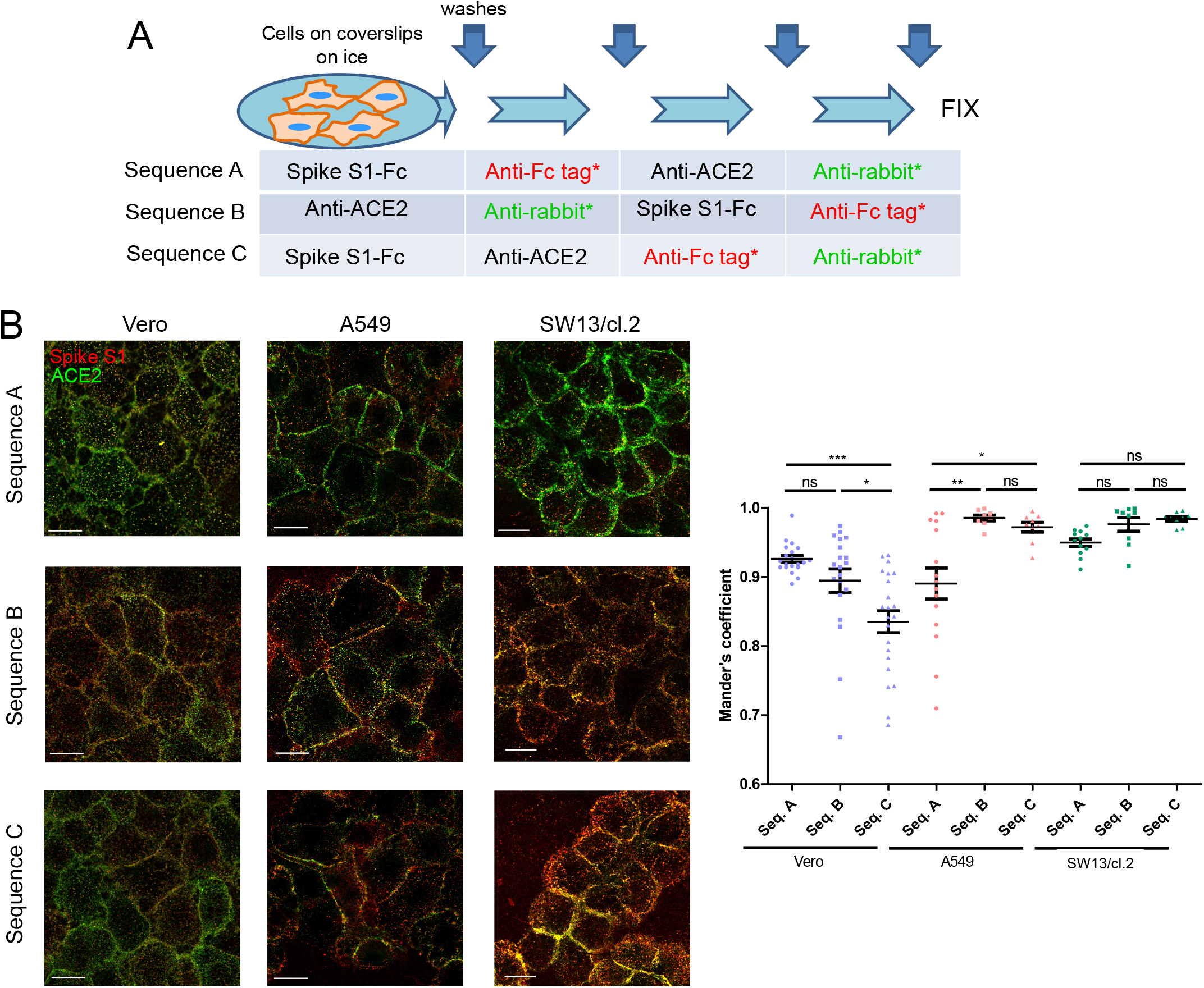
Detection of Spike constructs and ACE2 in several cell types employing different sequences for immunodetection. (A) Scheme of the incubation and washing steps performed for immunodetection. Incubations were carried out for 1 h in the cold. After each incubation coverslips were washed three times with 200 μl of cold PBS. At the end of the procedure, an additional washing step with water was performed before coverslips were allowed to dry and and mounted. (B) Representative images from the detection of Spike constructs and ACE2 in the indicated cell lines employing the different immunodetection sequences. The graph shows the colocalization between Spike S1-Fc and ACE2 fluorescent signals for every cell type and immunodetection sequence assayed. Colocalization is expressed as the percentage of Spike S1 colocalizing with ACE2 signal (Mander’s coefficient), measured applying the automatic Costes’ threshold (n≥8 per condition). Results are shown as mean values ± SEM; ns, non-significant, p > 0.05; *, p ≤ 0.05; **, p ≤ 0.01; ***, p ≤ 0.001 by ANOVA with Tukey’s post-test. Bars, 20 μm.

### Detection of ACE2 and vimentin on the surface of cells

In an earlier report, vimentin was identified as one of the proteins coimmunoprecipitating with ACE2 in Vero E6 cells exposed to SARS-CoV [23]. Therefore, we explored the presence of both proteins on the surface of A549 cells by immunofluorescence (Fig. 9A). For these assays, cells were incubated with a combination of the two primary antibodies followed by the secondary antibodies. A moderate degree of colocalization between vimentin and ACE2 immunoreactive signals was observed (Pearson’s correlation 0.36; overlap coefficient, 0.64; colocalization rate, 20%). Interestingly, colocalization was especially consistent at some intercellular segments as well as at certain spots, as highlighted in the colocalization mask (Fig. 9A). Remarkably, some of the colocalization spots appeared close to the nucleus and spanned multiple sections of the cell, as observed in orthogonal projections, being still noticeable at the upper cell layers (Fig. 9A), suggesting that both proteins colocalized at elongated structures. Interestingly, ACE2 has been recently shown to localize at the motile cilia of airway epithelial cells [52], which have been directly implicated in SARS virus infection [52, 53], as well as at the primary cilium of a kidney cell line [52]. While searching the literature for images of these structures, we spotted an intriguing overlap between the signal of vimentin, used as a fibroblast marker, and that of acetylated tubulin, a known marker of cilia [54, 55], in a study on cardiac fibrosis [56]. This prompted us to monitor the presence of ACE2 and vimentin at primary cilia in A549 cells. For this, we used several cell culture conditions, sample preparation strategies, including gentle washing to prevent deciliation, and cilia markers.

**Figure 9.**
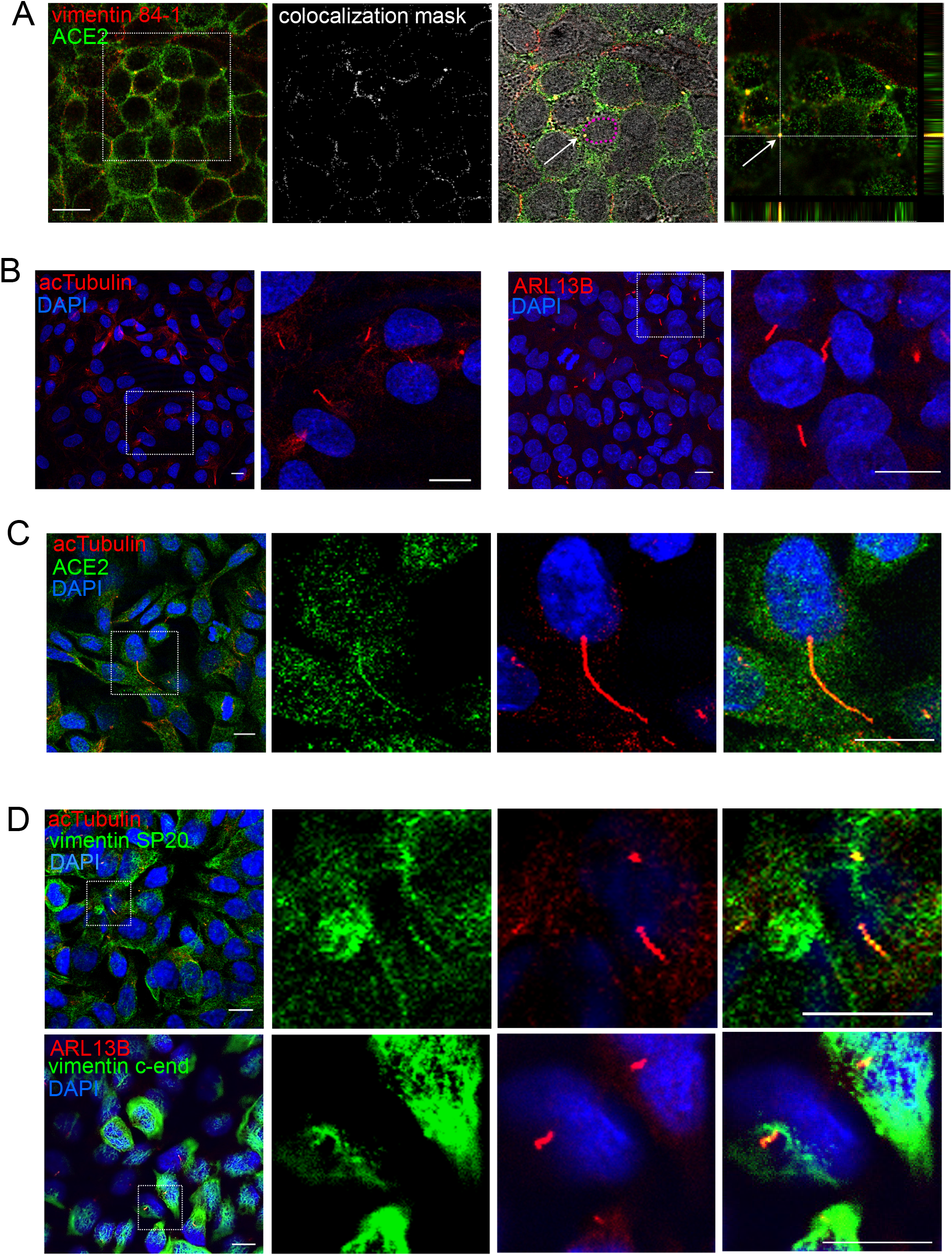
Detection of vimentin and ACE2 in A549 cells. (A) Live A549 cells were incubated simultaneously with anti-vimentin and anti-ACE2 antibodies, after which they were incubated with a combination of the corresponding secondary antibodies prior to fixation. An illustrative image is depicted showing points of colocalization at intercellular contacts at mid-cell height, highlighted in the colocalization mask. In addition, an overlay of the region of interest delimited by the dotted square with the bright field image is shown to illustrate the juxtanuclear position of one of the colocalization points (white arrow); the contour of the nucleus is highlighted in magenta. The far right panel shows the top section of the region of interest along with the orthogonal projections centered in the colocalization point marked with the arrow. Bars, 20 μm. (B) A549 cells were fixed with 4% (w/v) PFA and permeabilized with 0.1% (v/v) Triton for 5 min for staining with monoclonal antibodies against acetylated tubulin (acTubulin, left panels) or ARL13B (right panels). In each case, the regions of interest are enlarged at the right. Nuclei were stained with DAPI. (C) A549 cells fixed and permeabilized as in (B) were stained with anti-ACE2 and anti-acetylated tubulin antibodies. Images shown are single sections. (D) A549 cells were fixed with 2% (w/v) PFA, permeabilized with 0.1% (v/v) Triton X-100 and stained with anti-vimentin antibodies (SP20 or C-end) and either anti-acetylated tubulin or anti-ARL13B antibodies, as indicated. Images in (B), (C) and (D) are single sections. Bars, 10 μm.

Primary cilia could be distinguished in the A549 cell population based on their positive signal for acetylated tubulin, elongated appearance and presence as a single structure per cell, frequently stemming from a juxtanuclear position and being detectable at the upper cell layers (Fig. 9B). Cilia also showed a positive signal for ADP-ribosylation factor-like protein 13B (ARL13B), another known cilia marker (Fig. 9B). Importantly, cilia showed frequent ACE2 positive staining (Fig. 9C). Interestingly, the presence of vimentin immunoreactivity at ACE2 and acetylated tubulin or ARL13B-positive structures with the characteristics of primary cilia could be confirmed with several anti-vimentin antibodies including 84-1, SP20 and C-end, both in permeabilized (Fig. 9D) and non-permeabilized cells (Fig. 9A, 10A and C). As a negative control for these observations we studied vimentin-deficient SW13/cl.2 adrenocortical carcinoma cells. Although cells and from the cortex of the adrenal gland display imperfect primary cilia formation in relation with low levels of acetylated tubulin and ARL13B [57], we were able to spot some short structures positive for ACE2 and acetylated tubulin or ARL13B in SW13/cl.2 cells, consistent with short cilia (Suppl. Fig. 7). However, these structures did not show vimentin staining. Taken together, these results clearly indicate the presence of vimentin at the primary cilium in close proximity to ACE2 in A549 cells.

### Detection of Spike S1, ACE2 and vimentin at the primary cilia of A549 cells

Finally, we assessed whether the Spike S1 protein bound to primary cilia (Fig. 10). For these experiments a modification of sequence C was used. A549 cells were incubated with Spike S1-Fc prior to immunodetection with a combination of anti-ACE2 and anti-vimentin antibodies, followed by the corresponding secondary antibodies and, finally, with anti-human IgG for detection of the Fc-tag on the Spike construct, before fixation. Under these conditions, we confirmed that Spike S1 could be detected at certain spots at the cell periphery in close proximity to ACE2 and vimentin signals (Fig. 10A). However, spotting primary cilia on the sole basis of ACE2 immunoreactivity was not straight-forward. Therefore, we carried out complementary assays for staining of Spike S1-ACE2-acetylated tubulin (Fig. 10B) and Spike S1-vimentin-ARL13B (Fig. 10C). Both strategies allowed detection of Spike S1 on cilia in the proximity of ACE2 or vimentin-positive signals, and the corresponding cilia markers. Moreover, image analysis of these conditions with Leica software confirmed the colocalization of ACE2 and Spike signals (Pearson coefficient 0.6), and the moderate, but clear coincidence of vimentin and Spike S1 signals (Pearson coefficient of approximately 0.4), as illustrated by the colocalization masks shown in Fig. 10 (right panels). Taken together, these observations reveal the selectivity of the coincidence of Spike S1 and ACE2 or vimentin signals at primary cilia.

**Figure 10.**
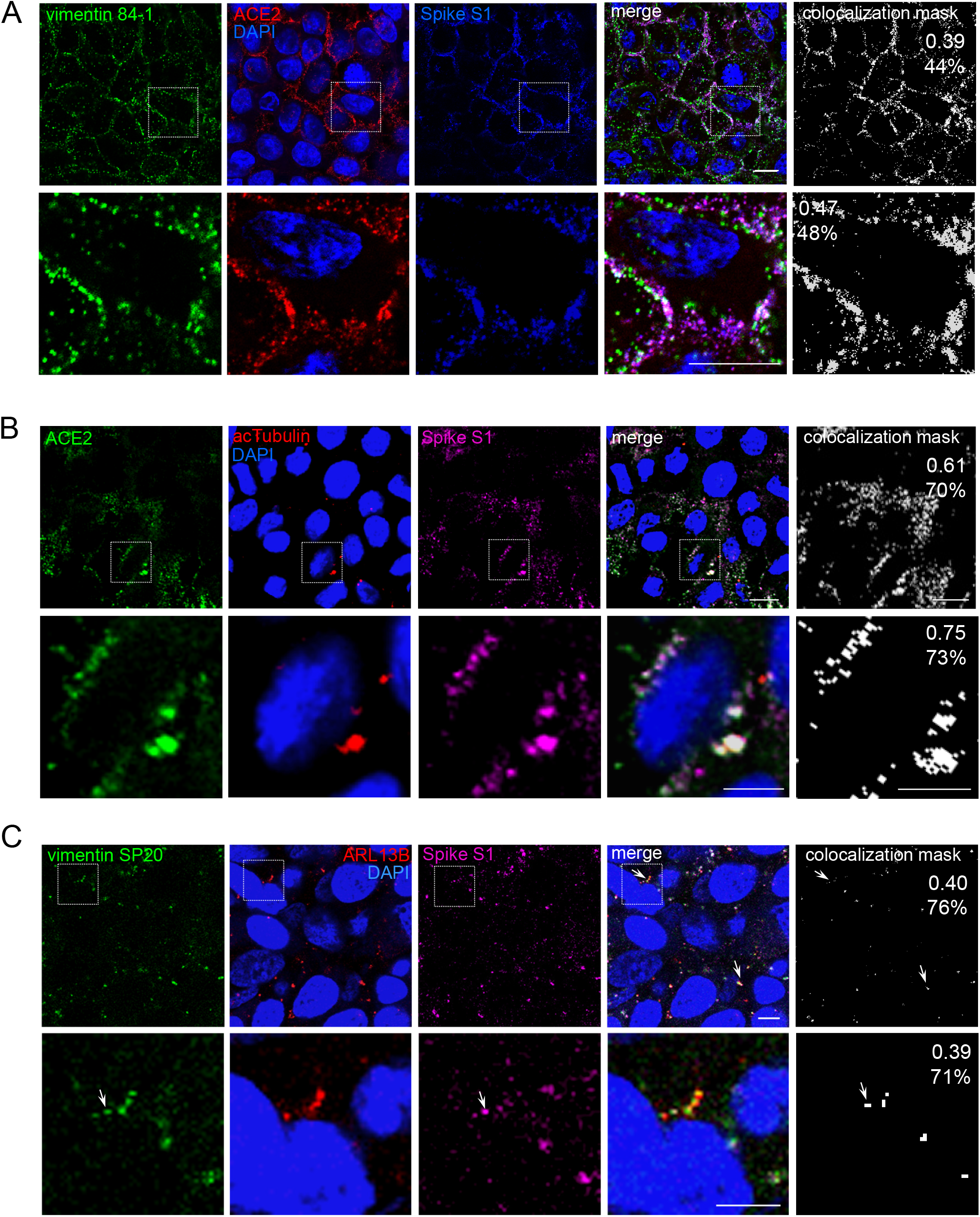
Detection of Spike S1, ACE2 and vimentin on cilia/acetylated tubulin-positive structures. A549 cells were incubated with Spike S1-Fc, for 90 min at 37ºC in complete medium. Non-permeabilized cells were stained with (A) anti-ACE2 and anti-vimentin 84-1 antibodies, (B) anti-ACE2 and anti-acetylated tubulin or (C) anti-vimentin SP20 and anti-ARL13B antibodies, followed by the corresponding secondary antibodies, and finally with anti-human IgG antibody to detect the Spike S1 Fc tag, before fixation. In (A) and (B) cells were fixed with 4% (w/v) PFA and in (C) with 2% (w/v) PFA. Images shown are single sections for each channel obtained by confocal microscopy and the corresponding overlays. Regions of interest are enlarged in the lower panels for each condition. Colocalization masks for the signals of (A) vimentin and Spike, (B), ACE2 and Spike, and (C) vimentin and Spike, are shown at the right as white signals on black background; numbers in insets correspond to the Pearson’s coefficient and the percentage of colocalization for the regions shown. Co-localization analysis was performed with Leica software. Arrows in C point to structures showing colocalization. Bars, 10 μm.

## Discussion

The docking of viruses onto cells is a multifaceted process in which multiple viral structures interact with various cellular components with diverse affinities to promote or facilitate viral entry. These interactions often involve cellular elements that cannot be considered true viral receptors since they cannot initiate viral RNA uncoating in the cytoplasm necessary for virus replication [58]. Regarding the interaction of SARS-CoV-2 with cells, numerous membrane-associated structures have been identified that can contribute to viral invasion, of which ACE2 is the most thoroughly studied. Moreover, involvement of other proteins as receptors or co-receptors for the virus, including the intermediate filament protein vimentin, has been hypothesized [12, 59, 60]. Taken together, the immunodetection studies reported here indicate a limited overall colocalization of vimentin with Spike proteins in non-permeabilized cells, but a consistent coincidence of vimentin and ACE2 at certain locations, especially at the primary cilium, which also appears to be a site for the attachment of SARS-CoV-2 Spike proteins. These observations open new possibilities to study a potential role of vimentin in the modulation of these structures and its connections with ACE2 function and viral infection.

In this study we have employed widely available tools, such as recombinant Spike protein constructs and several commonly used cell lines, to make an initial assessment of the presence of vimentin at the cell surface in relation to Spike binding sites. In our models, tagged Spike proteins lined the surface of several cell types yielding a rather homogeneous dotted pattern. Lipid rafts have been proposed as important points for viral entry and potential therapeutic targets [61]. Cholesterol supplementation has been reported to provoke the clustering of ACE2 and other viral entry factors at membrane sites with the characteristics of lipid rafts, thus increasing the potential binding sites for the virus [38], whereas a protective effect of strategies aimed at lowering or disrupting membrane cholesterol against coronavirus infections has been put forward [61, 62]. In addition, molecular dynamics simulations have proposed the existence of a ganglioside binding surface at the N-terminal domain of the SARS-CoV-2 Spike protein [63]. Nevertheless, the presence of ACE2 in lipid rafts is still controversial [64] and evidence on the association of SARS-CoV-2 with lipid rafts is mainly indirect. Under our conditions, the degree of colocalization of Spike protein constructs with CTXB as a marker of lipid rafts was low and did not support a preferential localization of Spike proteins in these membrane domains. Nevertheless, these results do not exclude the possibility that the virus may interact with lipid rafts in vivo.

Our next objective has been the detection of cell surface vimentin, which turned out not a straight forward task. The molecular form(s) of cell surface vimentin are still not thoroughly characterized (reviewed in [12, 65]). It appears that cell surface vimentin can exist in particular non-filamentous assembly states and/or bear certain posttranslational modifications [14, 66, 67]. Depending on the experimental system, phosphorylation, lipoxidation and/or oxidative modifications have been associated with the presence or increased cell surface exposure or secretion of vimentin [14, 46, 67]. Moreover, the accessibility of vimentin epitopes and the domains involved in the interaction with its partners in the extracellular medium may be variable and have not been always characterized in detail. In addition, fixation procedures, as shown herein, can either permeabilize or cause cell membrane damage allowing antibody entry and staining of the abundant cytoplasmic vimentin. This, in turn, can be the source of false positives above all in approaches that do not provide imaging, such as ELISA or flow cytometry. Therefore, detection and characterization of cell surface vimentin is a challenging task that requires working with non-permeabilized cells, and ideally non-fixed cells. Published images of vimentin detected at the cell surface do not respond to a universal pattern, and diverse arrangements including accumulations [68, 69], cell-membrane associated patches of variable extension [48, 70, 71], dotted patterns [48] and isolated dots [72] have been reported. Importantly, detection of cell surface vimentin relies on the availability of specific antibodies that perform adequately under the experimental conditions needed, i.e. absence of permeabilization or internalization, and do not show crossreactivity with other antigens in cells or reagents in the protocol.

Antibody-based detection of cell surface antigens is widely used although it is not exempt from limitations. Many of the antibodies employed herein have been validated by various methods (see Suppl. Table 1), including the confirmation of lack of signal in vimentin-deficient or knockout cells. Nevertheless, most of them have not been validated for the specific detection of cell surface vimentin. Under our conditions, incubation of live, non-permeabilized cells with a variety of anti-vimentin antibodies gave a dotted pattern non-homogeneously distributed along the cell surface. In addition, the presence of adhered vimentin or of focal membrane permeabilization contributed to the heterogeneity of the pattern of cell surface vimentin staining. Of the antibodies used, the 84-1 clone gave the strongest signal, followed by the V9 antibody in cells subsequently fixed with 4% (w/v) PFA. Nevertheless, controls of specificity carried out in two vimentin-deficient cell lines, namely SW13/cl.2 adrenal carcinoma and HAP1 (vim -), with either the V9 or the 84-1 antibodies still yielded a non-negligible background. A putative source for this background could be vimentin present in serum, e.g., in exosomes [73]; however, it was not abolished by culturing SW13/cl.2 cells in the absence of serum.

Alternatively, the possibility of a minor crossreactivity of the antibodies with other cellular structures needs to be considered. Indeed, certain anti-vimentin antibodies generated in patients show crossreactivity with some antigens from pathogens, such as streptococcal Hsp70 [74]. Moreover, certain vimentin epitopes can display similarities, at least in primary sequence, with other proteins. A BLAST search of the peptide ^106^ELQELNDR^113^, likely representing the epitope for 84-1 [49], retrieves hits in other intermediate filament proteins, including lamin, peripherin and desmin, as well partial hits in other proteins, such as the zinc finger protein ZC3H15. In turn, the V9 epitope shows partial homology to less related proteins, including nucleoporin, nuclear pore complex protein Nup153 isoform X2, and the immunoglobulin heavy chain junction region. These data raise the need of employing several antibodies to confirm the results and minimize the possibility of non-specific reactions. Nevertheless, neither the 84-1 nor the V9 antibodies recognize any peptide in lysates from vimentin-deficient cells analyzed by denaturing SDS-PAGE and western blot. Importantly, of the two monoclonal antibodies, 84-1 was the one giving a cleaner signal in dot blot when assayed against recombinant vimentin, Spike proteins and secondary antibodies, and hence became the preferred antibody in this study.

Remarkably, the overall colocalization of the vimentin immunoreactive signal with that of Spike protein constructs varied depending on the immunodetection sequence, being higher when Spike was detected in the first place (sequence A in Fig. 7). This phenomenon could be due to technical or mechanistic reasons. A high colocalization signal could be due to crossreactivity with the tagged Spike proteins employed or with neoepitopes or immune complexes formed during immunodetection. On the other hand, prior incubation with Spike and anti-human immunoglobulins could affect cells leading to higher vimentin exposure, although this may seem unlikely at 4ºC, or to focal cell damage. Otherwise, prior incubation with anti-vimentin antibodies could shield some Spike docking sites leading to alternative binding. In this context, a recent preprint reported the *in vitro* interaction between Spike-containing viral-like particles and vimentin, together with a blocking effect of some anti-vimentin antibodies on their uptake in an ACE2-overexpressing cell line [60]. Nevertheless, under our conditions, neither the lateral areas of cells with more intense vimentin signal nor the vimentin fragments adhered to the cell surface showed any particular enrichment of Spike binding.

Immunofluorescence of intact cells with anti-ACE-2 antibodies resulted in a robust punctate staining at the cell periphery in all cell types, consistent with previous reports [75]. This pattern was similar in several cell types, including A549 cells. Interestingly, levels and/or localization of ACE2 in A549 cells have been reported to vary depending on the proliferative state of cells, with postconfluent, less proliferative cells displaying higher ACE2 protein levels [43]. Here, we also observed a different ACE2 staining pattern in cells depending on the degree of confluency. Nevertheless, our results show a marked colocalization of ACE2 signals with the Spike proteins under the various incubation conditions.

Remarkably, ACE2 and vimentin immunoreactive signals displayed an apparently specific colocalization pattern, with coincidence of both signals at certain intercellular patches and distinct cellular structures compatible with primary cilia. The primary cilium is a structure that projects from the surface of most mammalian cells and plays key roles in the interaction with the environment and cell cycle control [76]. The primary cilium possesses a core of nine microtubules and a highly complex protein composition [77]. In turn, motile cilia are present at the apical surface of ciliated cells in the airways and other ducts. Notably, motile ciliated cells in the airways have been reported to originate from primary ciliated cells [78]. Interestingly, a recent article has reported the presence of ACE2 both at the motile cilia of epithelial airway cells and at the primary cilia of a ciliated kidney epithelial cell line [52]. Curiously, we observed that both ACE2 and vimentin-positive structures in the primary cilium were accessible in non-permeabilized cells in cold, while fixation and permeabilization allowed their detection in longer and more numerous cilia. Although additional studies in primary cells and tissue specimens need to be carried out to confirm these results, vimentin also seems to coincide with the signal of acetylated tubulin in images from primary rat cardiac fibroblasts and human tissue depicted in a recent report [56]. Some previously known connections between intermediate filaments and primary cilium include the regulatory role of several intermediate filament-associated proteins, including trichoplein and filamin A in ciliogenesis [79, 80], as well as the presence of a net of intermediate filaments around basal bodies [81]. Nevertheless, to the best of our knowledge, the participation of vimentin in cilium architecture had not been underscored.

The presence of ACE2 at cilia has been related to the involvement of these structures as primary points of SARS-CoV-2 docking for entry into cells. Indeed, in a study on human tissue, ciliated cells have been identified as selective targets for SARS-CoV-2 infection [52]. This is consistent with findings from other coronaviruses [53, 82], for which the presence of SARS-CoV structural proteins and viral particles associated to cilia was observed by electron microscopy [53]. Interestingly, SARS-CoV-2 infection has been proposed to associate with deciliation of invaded cells and cilia and flagellar dysfunction, which could occur through multiple mechanisms given the numerous interactions occurring between viral and cilia proteins (reviewed in [83, 84]). Indeed, a recent hypothesis envisages that many of the pathological alterations in COVID-19 could be the result of cilia dysfunction [83]. In this context, although the function of ACE2 in cilia is not known, a potential interference of SARS-CoV-2 with cilia functions through interaction with ACE2 has been speculated [52].

## Concluding remarks

The presence of vimentin has been reported to influence cell susceptibility to bacterial and viral pathogens, either in a positive or in a negative way, through various mechanisms, including behaving as an attachment factor at the cell membrane or influencing the replicative cycle [12, 20, 25, 85, 86]. Based on previous work on other coronaviruses, the potential role of vimentin in SARS-CoV-2 infection is currently a topic of interest [12, 59, 60], although the experimental evidence is so far limited. Our immunodetection studies in cell culture models incubated with Spike protein constructs point to a scarce overall colocalization of vimentin with the signals from Spike and ACE2. Nevertheless, they highlight the presence of vimentin at selective cellular structures, namely primary cilia, where these proteins appear to be enriched. It should be noted that colocalization as described herein indicates proximity of the proteins detected but does not imply a direct interaction. Therefore, additional studies will be performed to explore this possibility. Nevertheless, these observations call for a detailed characterization of the presence of vimentin in primary cilia, as well as of its potential role in interaction with pathogens at this location.

## Acknowledgements

This work was supported by grants CSIC PTI Global Health (PIE 202020E223/CSIC-COV19-100), RTI2018-097624-B-I00 from Micinn (Agencia Estatal de Investigación), Spain and ERDF. Á.V.P. and P.G.J. are the recipients of predoctoral contracts BES-2016-076965 and PRE2019-088194, respectively, from Micinn, Spain.

## Abbreviations

ACE2: Angiotensin converting enzyme 2
ARL13B: ADP-ribosylation factor-like protein 13B
CTXB: cholera toxin B subunit
TMPRSS2: transmembrane protease serine 2

