## Supplementary information for "Immunolocalization studies of vimentin and ACE2 on the surface of cells exposed to SARS-CoV-2 Spike proteins"

Vasiliki Lalioti, Silvia González-Sanz, Irene Lois-Bermejo, Patricia González-Jiménez, Álvaro Viedma-Poyatos, Andrea Merino, María A. Pajares, Dolores Pérez-Sala

Department of Structural and Chemical Biology. Centro de Investigaciones Biológicas Margarita Salas, CSIC. 28040 Madrid, Spain

Short title: Cell Surface detection of vimentin, ACE2 and Spike

Please address correspondence to:

Dolores Pérez-Sala
Centro de Investigaciones Biológicas Margarita Salas, CSIC
Ramiro de Maeztu, 9
28040 Madrid, Spain


Supplementary figures

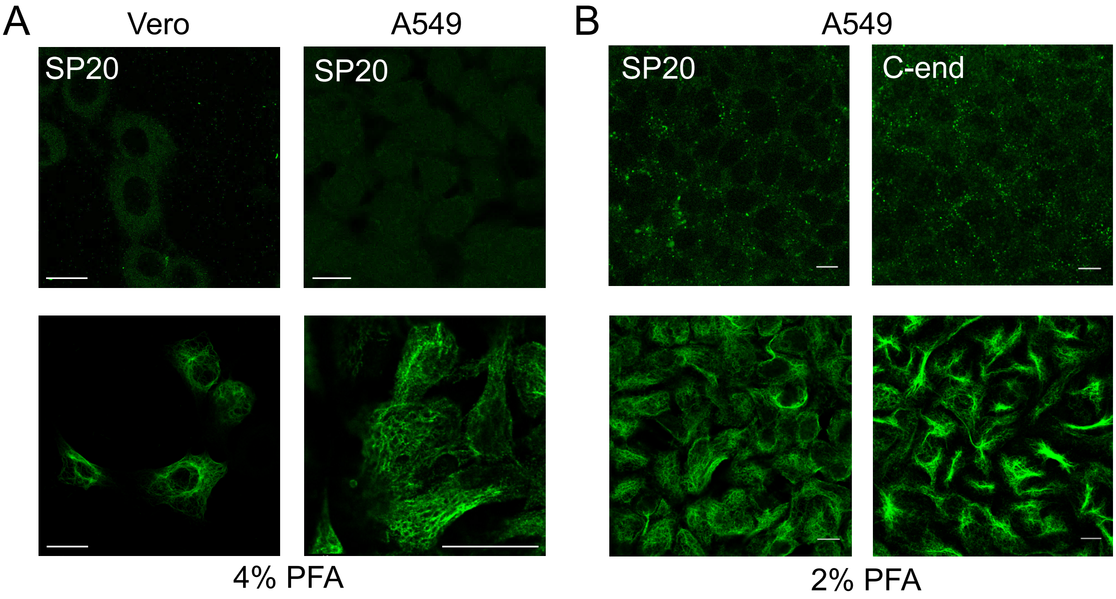

Supplementary Figure 1. Immunodetection of vimentin using two fixation procedures. (A) Vero or A549 cells were incubated with anti-vimentin SP20 rabbit monoclonal antibody at 1:200 dilution for 1 h in the cold, followed by incubation with Alexa-488-conjugated anti-rabbit immunoglobulins at 1:200, before fixation with 4% (w/v) PFA in the cold (upper panels), or after fixation and permeabilization (lower panels). Representative images at mid cell height are shown. Bars, 20 μm. (B) A549 cells were incubated with the indicated anti-vimentin antibodies, and the corresponding secondary antibodies, before fixation with 2% (w/v) PFA (upper panels) or after fixation and permeabilization (lower panels) for detection of cytoplasmic vimentin. Bars, 10 μm.

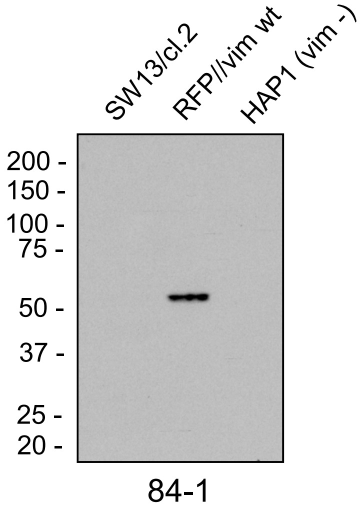

Supplementary Figure 2. Immunoblot analysis of vimentin in several cell types. Total cell lysates from the indicated cell types containing 30 μg of protein were analyzed by SDS-PAGE followed by immunoblot with the 84-1 anti-vimentin antibody. SW13/cl.2 cells stably transfected with the bicistronic vector RFP//vimentin wt as detailed in Materials and Methods (RFP//vim wt) were used as positive control.

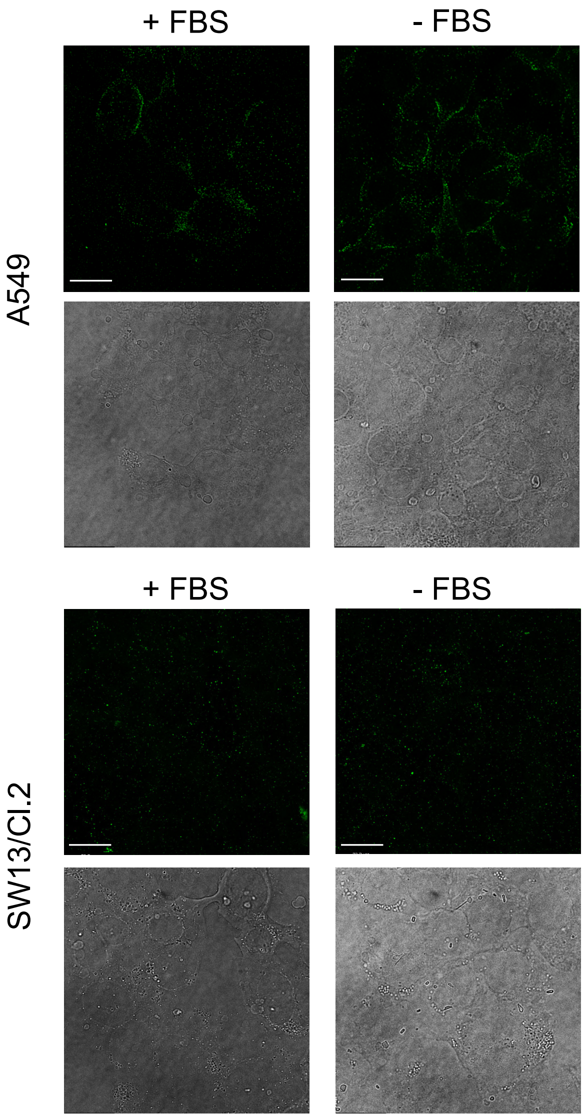

Supplementary Figure 3. Effect of serum depletion on the detection of vimentin at the cell surface. A549 or SW13/cl.2 cells were cultured in the presence or absence of FBS for 24 h before immunodetection of vimentin in live cells with the 84-1 monoclonal antibody. Images shown are fluorescent confocal single sections at mid-cell height (upper panels) and bright field images (lower panels). Bars, 20 µm.

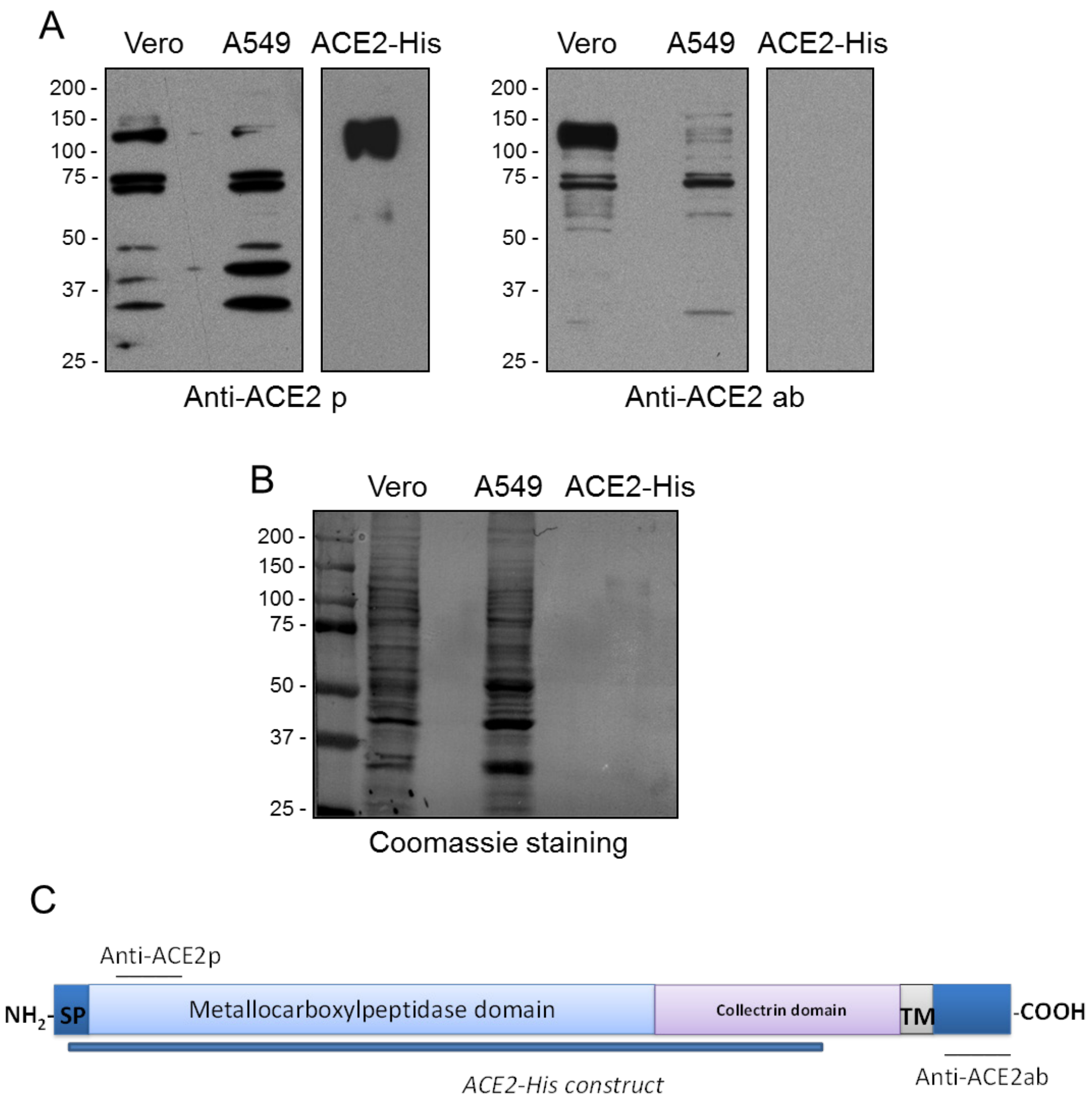

Supplementary Figure 4. Immunoblot analysis of ACE2 in several cell types. (A) Total cell lysates from the indicated cell types containing 30 μg of protein, or 100 ng of purified ACE2 protein were analyzed by SDS-PAGE followed by immunoblot with the indicated anti-ACE2 antibodies. (B) Total protein on blots was visualized by staining with Simply Blue. (C) Scheme depicting the sequence of ACE2, the location of the epitopes for the antibodies used and the region spanned by the recombinant protein. SP, signal peptide; TM, transmembrane domain.

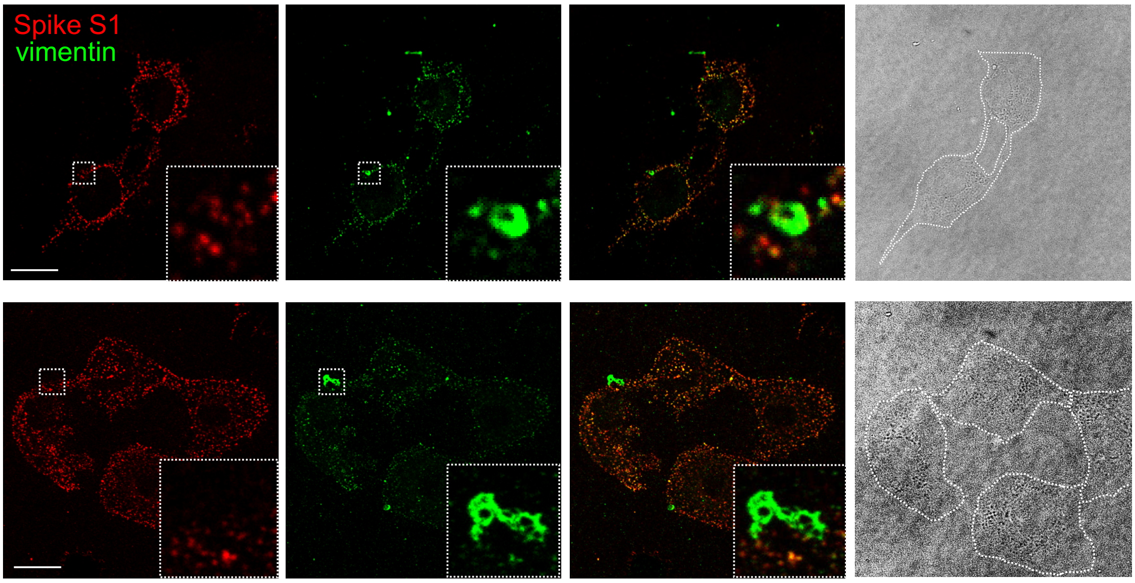

Supplementary Figure 5. Detection of Spike binding and vimentin in cells showing vimentin adhered to the cell surface. Vero cells were incubated with Spike-S1-Fc prior to immunodetection of vimentin following the protocol outlined in Figure 6 as sequence A. Single sections of the individual channels at mid cell height as well as the merged images are shown. Insets show enlarged views of the regions of interest marked by dotted squares. Panels on the right depict bright field images with the cell contours outlined. Bars, 20 µm.

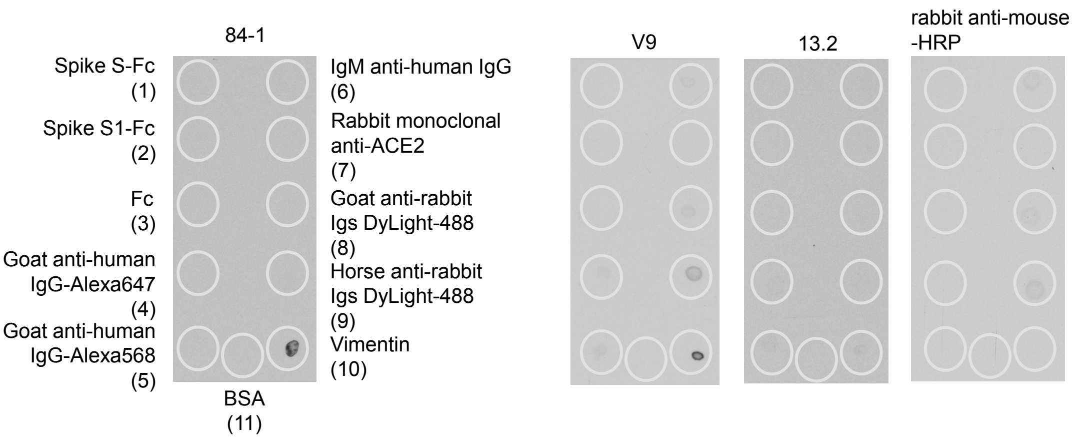

Supplementary Figure 6. Dot blot of several protein constructs and antibodies. Five µl of the indicated reagents at the concentrations used for immunodetection were spotted on nitrocellulose membrane at the positions denoted by the grey circles and allowed to dry before blocking with 2% (w/v) non-fat powdered milk in T-TBS, followed by incubation with the indicated anti-vimentin antibodies at 1:500 dilution in 1% (w/v) BSA in T-TBS, and HRP-conjugated rabbit anti-mouse immunoglobulins for detection with ECL. The HRP-conjugated antibody alone was used as a control. The amounts of protein spotted in each case were as follows: Spike S-Fc, Spike S1-Fc, Fc, goat anti-human IgG-Alexa647, goat anti-human IgG-Alexa568 and BSA, 0.05 µg; IgM anti-human IgG and rabbit monoclonal anti-ACE2, 0.025 µg; goat anti-rabbit Igs DyLight-488 and horse anti-rabbit Igs DyLight-488, 0.035 µg; and vimentin 0.035 µg. Note that the 84-1 antibody only detected vimentin. The V9 antibody gave a detectable background with the anti-rabbit antibodies, spots 8 and 9, and a fainter signal with the goat anti-human IgG antibodies, spots 4 and 5. The background with anti-rabbit antibodies could be due in part to binding of the HRP-conjugated rabbit anti-mouse antibody used as secondary antibody for detection. The 13.2 antibody gave a very faint signal with recombinant vimentin.

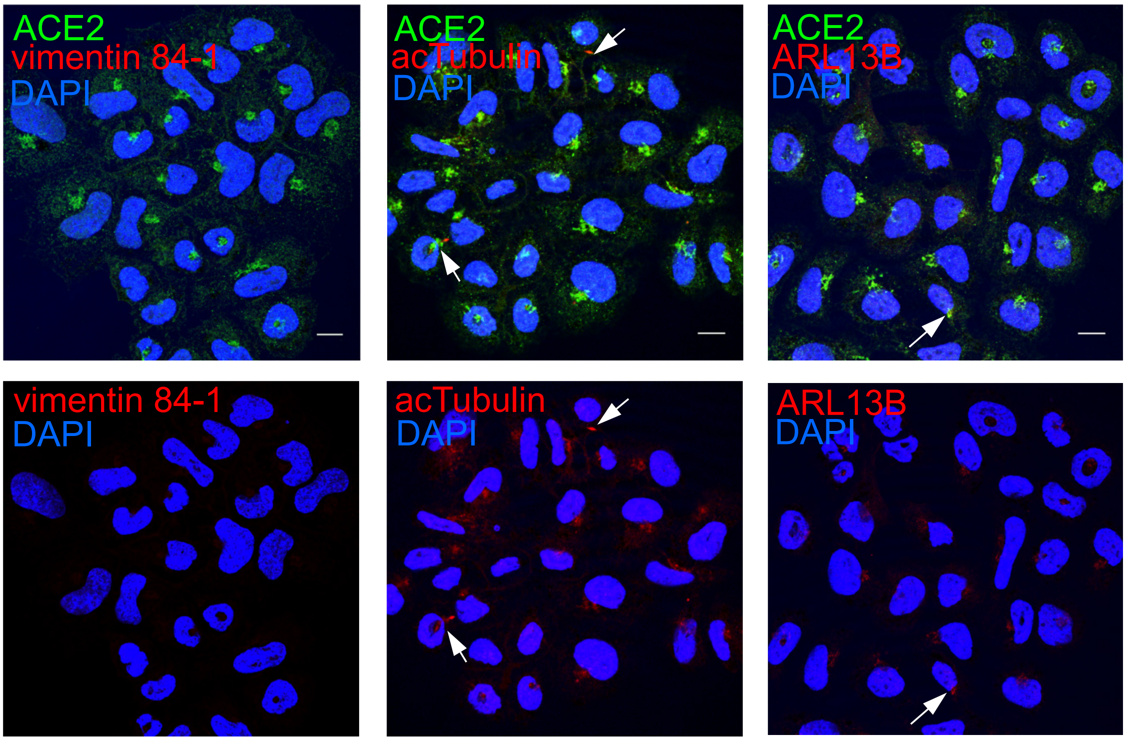

Supplementary Figure 7. Detection of vimentin and ACE2 in combination with markers of primary cilia in SW13/cl.2 cells. Vimentin-deficient SW13/cl.2 cells were fixed with 4% (w/v) PFA, permeabilized with 0.1% (v/v) Triton and stained with the indicated antibodies. Upper panels show merge images of all channels and lower panels show staining with the individual anti-vimentin, anti-acetylated tubulin or anti-ARL13B antibodies. Nuclei were counterstained with DAPI. White arrows mark incipient structures positive for acetylated tubulin or ARL13B immunoreactivity. Bars, 10 μm.

| Antibodies | | | | | | | |
|---|---|---|---|---|---|---|---|
| **Antigen** | **Name** | **Type** | **Immunogen** | **Epitope** | **Vendor** | **Reference** | **Knockout validated** |
| **Primary antbodies** | | | | | | | |
| vimentin | V9 | mouse monoclonal | purified porcine vimentin | C-terminal | Santa Cruz Biotech. | sc-6260 | Yes |
| | 13.2 | mouse monoclonal | human foreskin fibroblast extract | Not determined | Sigma | V5255 | |
| | 84-1 (CSV) | mouse monoclonal | human recombinant vimentin | Putative, residues 106-113 | Abnova | H00007431-M08J | |
| | SP20 | rabbit monoclonal | human recombinant vimentin | Not determined | Thermo | MA5-14564 | Yes |
| | C-end (vim 453-466) | goat polyclonal | peptide: QVINETSQHHDDLE | C-terminal end | Everest Biotech | EB11207 | |
| Ace2 | | rabbit polyclonal | synthetic peptide | N-terminus | Invitrogen | PA5-20045 | |
| | ACE2 788-805 | rabbit polyclonal | peptide: CKGENNPGFQNTDDVQTSF | C-terminus | Abcam | ab15348 | |
| Spike | SARS-CoV-2 (2019-nCoV)- | rabbit monoclonal | | Not determined | Sino Biological | 40150-R007 | |
| | Spike antibody clone 007 | | | | | | |
| ARL13B | C5 | mouse monoclonal | | C-terminal, residues 414-428 | Santa Cruz Biotech. | sc-515784 | |
| **Secondary antibodies** | | | | | | | |
| | Anti-human IgG (H+L)-Alexa fluor 568 | goat polyclonal | | | Invitrogen | A21090 | |
| | Anti-human IgG Fc-Dylight 488 | goat polyclonal | | | Invitrogen | SA5-10134 | |
| | Anti-human IgG Alexa 647 | goat polyclonal | | | Invitrogen | A21445 | |
| | Anti-human IgG (H+L) Alexa fluor 647 | alpaca polyclonal | | | Jackson Immun. Lab. Inc. | 609-605-213 | |
| | Anti-mouse F(ab')2 Alexa 488 | goat polyclonal | | | Jackson Immun. Lab. Inc. | 115-546-062 | |
| | Anti-mouse Igs Alexa 647 | goat polyclonal | | | Invitrogen | A21235 | |
| | Anti-rabbit Dylight 488 | horse polyclonal | | | Vector laboratories | DI-1088 | |
| | Anti-rabbit Alexa 647 | goat polyclonal | | | Invitrogen | A21244 | |

Supplementary Table 1. Antibodies used in this study.
